# Spatial Morphoproteomic Features Predict Uniqueness of Immune Microarchitectures and Responses in Lymphoid Follicles

**DOI:** 10.1101/2024.01.05.574186

**Authors:** Thomas Hu, Mayar Allam, Vikram Kaushik, Steven L. Goudy, Qin Xu, Pamela Mudd, Kalpana Manthiram, Ahmet F. Coskun

**Affiliations:** Wallace H. Coulter Department of Biomedical Engineering, Georgia Institute of Technology and Emory University, Atlanta, GA, USA; School of Electrical and Computer Engineering, Georgia Institute of Technology, Atlanta, GA, USA; Department of Otolaryngology–Head and Neck Surgery, Emory University School of Medicine, Atlanta, Georgia, U.S.A; Cell Signaling and Immunity Section, Laboratory of Immune System Biology (LISB), National Institute of Allergy and Infectious Diseases (NIAID), National Institutes of Health (NIH), Bethesda, MD, USA; Division of Pediatric Otolaryngology, Children’s National Hospital, Washington, DC, USA, Division of Otolaryngology, Department of Surgery, George Washington University School of Medicine and Health Sciences, Washington, DC, USA; Interdisciplinary Bioengineering Graduate Program, Georgia Institute of Technology, Atlanta, GA, USA; Winship Cancer Institute, Emory University, GA, USA; Parker H. Petit Institute for Bioengineering and Bioscience, Georgia Institute of Technology, 315 Ferst Dr. NW, Atlanta, GA 30332

## Abstract

Multiplex imaging technologies allow the characterization of single cells in their cellular environments. Understanding the organization of single cells within their microenvironment and quantifying disease-status related biomarkers is essential for multiplex datasets. Here we proposed SNOWFLAKE, a graph neural network framework pipeline for the prediction of disease-status from combined multiplex cell expression and morphology in human B-cell follicles. We applied SNOWFLAKE to a multiplex dataset related to COVID-19 infection in humans and showed better predictive power of the SNOWFLAKE pipeline compared to other machine learning and deep learning methods. Moreover, we combined morphological features inside graph edge features to utilize attribution methods for extracting disease-relevant motifs from single-cell spatial graphs. The underlying subgraphs were further analyzed and associated with disease status across the dataset. We showed that SNOWFLAKE successfully extracted significant low dimensional embedding from subgraphs with a clear separation between disease status and helped characterize unique cellular interactions in the subgraphs. SNOWFLAKE is a generalizable pipeline for the analysis of multiplex imaging data modality by extracting disease-relevant subgraphs guided by graph-level prediction.

## Introduction

Snowflakes can form into various shapes and patterns depending on different environmental parameters including air temperature, air humidity, dust particles, etc. Depending on these parameters, snowflakes can form into plates, needles, solid prisms, or the most known shape, dendrites ^1^. Further, snowflakes form in stages where the vapor in the air can transform into the quasi-liquid phase and then to the solid phase to form the first snow crystal of the snowflake. This can initiate the formation of more crystals at the boundary sites to finally form complex shapes of snowflakes through thousands of these steps ^2^. The unique property of snowflakes attracted great attention and how the immune system uniquely responds to infections presents opportunities to draw parallels between snowflakes and the microstructural organization of immune organs in tissues.

Akin to the snowflakes, germinal centers are microstructures in lymphoid tissues that form into stages where the antigens can either exist in the form of soluble antigens or present through antigen-presenting cells (APCs) including follicular dendritic cells (FDCs), After the antigen encounter, B cells upregulate the expression of the chemokine receptor CCR7. This further drives B-cell migration towards ligands of CCR7 including CCL19 and CCL21, which exist at the T-cell and B-cell border of the primary follicles ^3^. Activated T cells which have upregulated the chemokine receptor CXCR5 also migrate to the T-cell B-cell border, where they have long-lived interactions with cognate antigen-presenting B cells. B cells then bind to CD4+ T-cells which further initiate B cell’s division and differentiation into either short-lived plasma cells or memory B cells or ultimately initiate a GC response ^4,5^ (**Supplementary Fig. 1a**). Moreover, primary follicles (including multiple types of B cells) and germinal centers (subsets of follicles with proliferating B cells undergoing somatic hypermutation) display unique features in their sizes, heterogeneous shapes and orientation depending on several factors. The presence of FDCs is considered to be the most critical component for follicular retention and organization ^6,7^. FDCs-ablated mice have disrupted primary follicle architecture with B-cells overlapping with the T-cells zone or the stromal network. This also results in unrestricted positioning of several cellular phenotypes including T cells and dendritic cells as well as chemokines gradients (ex: CCL21, CXCL13) and extracellular matrix networks (e.g., Collagen). Ablation of FDCs also affects the migration and clustering behavior of naive B-cells which, under normal conditions, helps maintain germinal center response. As a result, FDC-ablated mice have a complete loss of the GC response after immunization signifying the crucial role of FDCs in initiating the primary follicles and maintaining the GCs ^7^. Beyond biological mechanisms driving follicle shapes and cellular organization, we reasoned that the uniqueness of follicles and germinal centers would be preserved whether subjects are infected with SARS-CoV-2 viruses or uninfected (**Supplementary Fig. 1a**). Thus, computational approaches should be able to resolve follicle morphology and cell organization using emerging deep-learning techniques, including graph-based geometric learning methods.

Graph neural networks (GNN) have shown ground-breaking performance on many deep learning tasks ^8^. Modeling tumor microenvironment single-cell spatial organization as graphs and applying GNN have predicted response and survival at the patient level using various types of imaging modalities such as multiplex imaging^9,10^ and H&E ^11,12^. On the other hand, cell morphology analysis has been used to study the classification of cell phenotypes ^13,14^. However, there is a lack of study between tissue morphological structures and proteomics cellular data from multiplexed imaging of lymphoid tissues.

To incorporate multimodal features of morphology and cell networking, we propose SNOWFLAKE (Spatial siNgle-cell Organization With Formation LeArning and Knowledge Embedding), a multi-modal graph-based learning method combining spatial morpho-proteomic features (both follicle morphology and multiplexed proteomic markers of cell compositions) in a single pipeline within multiplexed proteomic images of lymphoid tissues (**Supplementary Fig. 1b**). SNOWFLAKE allows single-cell protein expression profiles combined with structure shape descriptors and single-cell position-aware graphs of follicles and germinal centers to predict immune response and quantify the uniqueness of tissue microenvironments. Moreover, SNOWFLAKE incorporates graph neural network interpretation methods such as gradient-based ^15^ or mutual information-based ^16^ to explain predictions from single-cell subgraphs that allow the extraction of single-cell graphlets biomarkers. Compared to existing methods utilizing graph neural networks for patient phenotype predictions from cellular-level graph analysis, SNOWFLAKE allows model explainability at the subgraph level while incorporating morphological features (**Supplementary Fig. 1b**).

## Results

### SNOWFLAKE workflow for spatial morphoproteomic features modeling

To demonstrate the ability of SNOWFLAKE to model spatial morpho proteomic features, we first constructed a large dataset library of various human tonsil and adenoid tissues from multiplex imaging technologies^17^ (**Supplementary Table 1 and Methods**). The SNOWFLAKE pipeline starts with extraction of single-cell proteomics, morphological and structural information from imaging data and then models single-cell neighboring information from multiplexing imaging as single-cell spatial graphs (**Fig. 1a**). Using the single-cell spatial graphs, SNOWFLAKE predicts the uniqueness of immune microarchitectures and responses in lymphoid follicles using spatial-morpho-proteomics features while providing subgraph explanation of important cell and their neighboring cells (**Fig. 1b**). For incorporating morphological features in our SNOWFLAKE pipeline, we used a PCA dimension reduction after shape alignment, sampling, and smoothing ^18^ (**Supplementary Fig. 2 and Methods**). We successfully transformed segmented boundaries of follicles and their corresponding germinal center into a low-dimensional representation that captures the morphological variation in our dataset **(Supplementary Fig. 3)**.

**Fig. 1.**
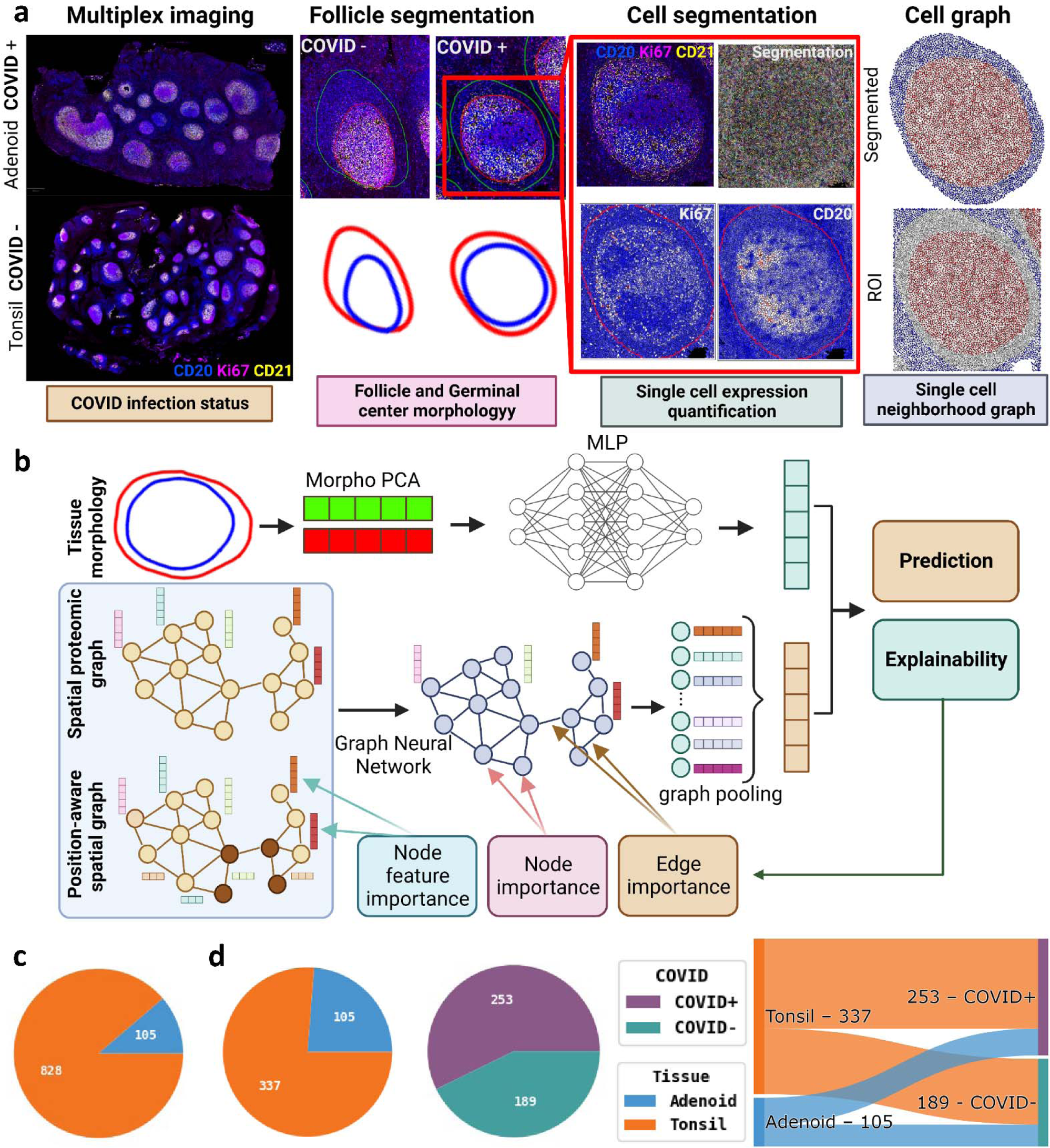
Overview of SNOWFLAKE. **a.** Raw data obtained from multiplex imaging showing CD20 (Blue), CD21 (yellow), and Ki67 (magenta) staining from tonsil and adenoid tissues with and without COVID infection. Follicles and corresponding germinal centers are segmented using multiplex panels (n=930 follicles and n=775 germinal centers). Single cells are segmented (n=8879749 cells), and cell neighborhood graphs are extracted based on cell spatial location. **b.** SNOWFLAKE prediction pipeline combining morphological PCA reduction of the follicle and germinal center shapes with the spatial proteomic graph for prediction of patient-level status. An explainability framework is used to understand the node, edge, and node feature importance for the prediction task. **c.** Pie graph showing the distribution of follicle database tissue distribution. **d.** Pie graph and Sankey plot showing the distribution of NIH-COVID follicle database tissue distribution, COVID status distribution.

The primary follicles and the germinal centers of human tonsil and adenoid tissues were manually annotated using CD20, CD21, CD3, and Ki67 signals therefore creating a diverse database of follicles (n=930). Next, using a published dataset with COVID-19 infection status of lymphoid tissues ^17^ from children, we curated a database of follicles COVID-19-convalescent children’ and uninfected controls and referred to it as NIH-COVID (**Table 1**, **Fig. 1c and Supplementary Fig. 4**). Children infected with SARS-CoV-2 undergo humoral and cellular immune responses in lymphoid tissues of the upper respiratory tract where infection and viral replication take place^17^. It has been shown that tonsils and adenoids retain SARS-CoV-2 RNA even months after the acute infection which leads to SARS-CoV-2-specific GC B cells in the tonsils and adenoids. In Adenoids, persistent enrichment of IgG+ and IgM+ GC B cells in convalescence is observed. Moreover, studies have shown prolonged affinity maturation of B cells in germinal centers months after infection from antigen persistence ^19–21^. It is therefore important to identify uniqueness around B-cell follicles from multiplex protein imaging. The NIH-COVID multiplex imaging provides a unique opportunity to study the spatial organization of single cells correlated with durable humoral immunity from SARS-CoV-2 infection. Furthermore, this approach could be extended to understand germinal center response following vaccination^22^, which is crucial for the development of immune memory and long-term protection against the virus.

**Table 1.**
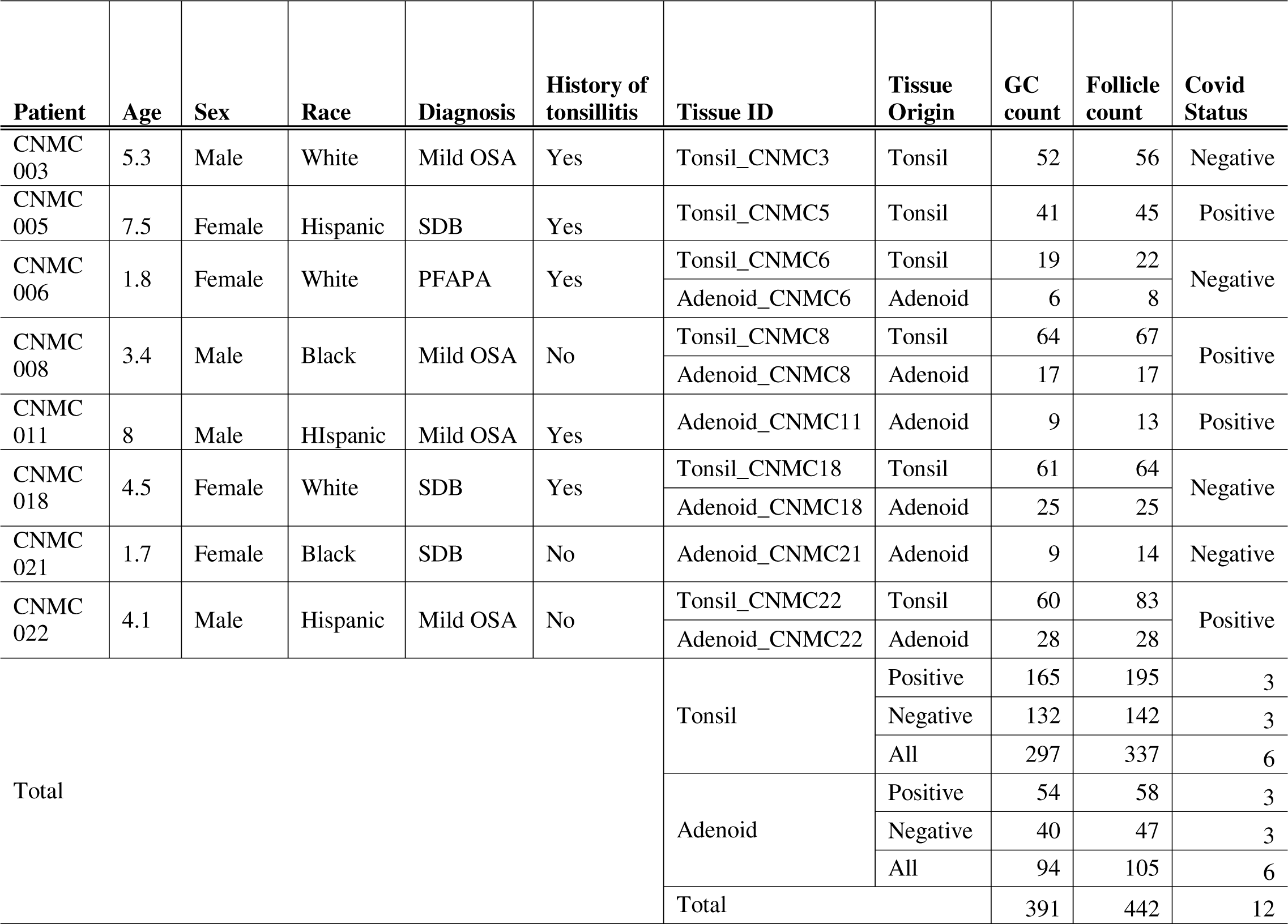
Statistical summary of the NIH-COVID dataset.

### SNOWFLAKE predicts COVID-19 infection status from lymphoid follicles

We applied the SNOWFLAKE pipeline to predict COVID-19 status in the NIH-COVID dataset from lymphoid follicles’ morphological (shape) and proteomic (antibody-based protein imaging) features. Single cells were segmented using multiplex markers in a semi-supervised fashion (**Methods**). We benchmarked SNOWFLAKE prediction with Machine Learning (ML) models on follicle-level mean expression, Multi-Layer Perception (MLP) models on follicle-level mean expression, and a Multi-Instance-Learning (MIL) MLP model on cell-level expressions (**Fig. 2a**). In the ML setting, we averaged the single-cell expression for all markers at the follicle level and tested the predictive power of mean expression only, morphological feature only, and concatenated features from mean expression and morphology. The MLP model is used similarly to the ML setting, where we used a multi-layer perception neural network for prediction (**Supplementary Table 2)**. In contrast, the MIL setting uses mean expressions at the single-cell level and uses a multi-layer perception network to learn cell-level embedding and a pooling layer is used to aggregate cell-level embedding for the follicle-level prediction. The graph neural network (GNN) pipeline is similar to the MIL while adding edge connectivity between cells by looking at the spatial neighboring information of single cells (**Methods**) therefore allowing aggregation of neighboring information in the feed-forward network. Moreover, we compared various Graph Neural Network (GNN) layers and graph pooling layers to obtain a comprehensive comparison between the different models (**Supplementary Table 3** and Methods). For studying the generalizability of our models with a relatively small cohort, we used a Monte Carlo 5-fold Cross-validation with a 30% validation set split to evaluate each model.

**Fig. 2.**
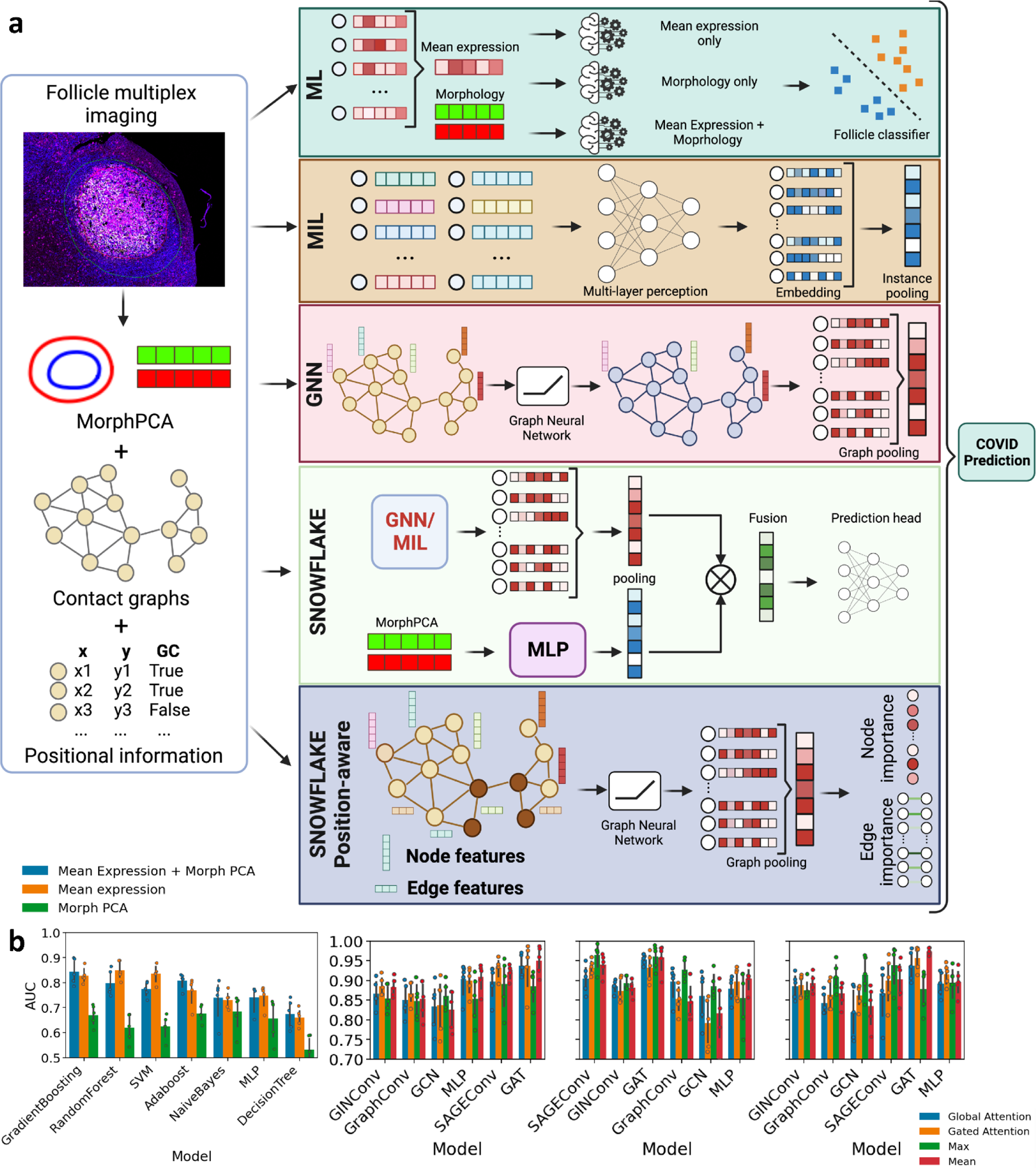
Overview of patient COVID status prediction pipeline. **a.** Follicle morphological features, single-cell spatial graphs, and positional information are extracted from follicle multiplex imaging. SNOWFLAKE pipeline and position-aware SNOWFLAKE pipeline are benchmarked against machine learning (ML) pipeline which considers only mean expression and morphological features. Multi-instance learning (MIL) framework is also compared by considering all single-cell expression levels across follicles without cell spatial information. Finally, the graph neural network (GNN) framework is compared by taking into account of simple cell neighborhood information for the prediction pipeline. SNOWFLAKE pipeline combines GNN pooling with Morph PCA pipeline using a fusion layer for prediction. Position-aware SNOWFLAKE takes into consideration the pairwise spatial distance and angle between each pair of connected cells in the spatial graph and allows node importance and edge importance downstream analysis from the model trained to predict COVID status. **b.** Bar plot showing the model AUC scores for diverse models in ML settings (left) with 5-fold cross-validation. Bar plot showing the model AUC scores for diverse graph neural network layers in the SNOWFLAKE pipeline (right) with 5-fold cross-validation.

For COVID-19 status prediction, SNOWFLAKE achieved a best model ROC AUC of 0.973 ± 0.009 compared to 0.915 ± 0.027 from MIL-MLP model, 0.748 ± 0.045 from MLP mean expression, and 0.85 ± 0.038 from ML mean expression (**Fig. 2b and Table 2**). This shows that combined spatial morpho-proteomic features from follicles in the SNOWFLAKE pipeline are indicative of COVID-19 infection status better than single-cell expression only. This indicates that COVID-19 infection is driving changes to the shape, size, cellular composition, and arrangement of the follicles.

**Table 2.** SNOWFLAKE model performance evaluation benchmarked with various model type.

| Type | Model | Pooling Strategy | Morphology Fusion | Accuracy | AUC | F1 |
| --- | --- | --- | --- | --- | --- | --- |
| Mean | Adaboost | | Mean Expression + Morph PCA | $0.767 \pm 0.043$ | $0.808 \pm 0.031$ | $0.476 \pm 0.062$ |
| | DecisionTree | | Mean Expression + Morph PCA | $0.74 \pm 0.049$ | $0.675 \pm 0.059$ | $0.497 \pm 0.07$ |
| | GradientBoosting | | Mean Expression + Morph PCA | $0.817 \pm 0.052$ | $0.844 \pm 0.051$ | $0.554 \pm 0.08$ |
| | NaiveBayes | | Mean Expression + Morph PCA | $0.663 \pm 0.051$ | $0.74 \pm 0.084$ | $0.504 \pm 0.07$ |
| | RandomForest | | Mean expression | $0.812 \pm 0.032$ | $0.85 \pm 0.038$ | $0.502 \pm 0.052$ |
| | SVM | | Mean expression | $0.806 \pm 0.028$ | $0.835 \pm 0.035$ | $0.387 \pm 0.067$ |
| | MLP | | Mean expression | $0.79 \pm 0.036$ | $0.748 \pm 0.045$ | $0.268 \pm 0.108$ |
| MIL | MLP | Gated Attention | Tensor Decomposition | $0.815 \pm 0.029$ | $0.915 \pm 0.027$ | $0.857 \pm 0.02$ |
| | GAT | Mean | Tensor Decomposition | $0.926 \pm 0.01$ | $0.973 \pm 0.009$ | $0.938 \pm 0.008$ |
| | GCN | Max | Tensor Decomposition | $0.827 \pm 0.025$ | $0.913 \pm 0.052$ | $0.856 \pm 0.024$ |
| | GINConv | Max | Expression Only | $0.795 \pm 0.025$ | $0.893 \pm 0.017$ | $0.82 \pm 0.028$ |
| | GraphConv | Max | Expression Only | $0.842 \pm 0.017$ | $0.927 \pm 0.026$ | $0.855 \pm 0.013$ |
| | SAGEConv | Max | Expression Only | $0.91 \pm 0.04$ | $0.964 \pm 0.033$ | $0.924 \pm 0.031$ |

The improvement of the ROC AUC metric for COVID-19 status prediction of single-cell expression in the MIL model compared to mean expression features when using ML models or MLP model indicates proteomic distribution inside the follicle is predictive of COVID-19 status compared to aggregation of all cell expression levels. The individual cell embedding combined with an aggregation layer for prediction better captures the heterogeneity of the follicles. Moreover, the GNN models improved the ROC AUC when compared to the MIL model showing the importance of single-cell spatial information for model prediction. It is interesting to see that the max pooling layer performed better across various graph neural network layers (Table 2) showing that a subset of neighboring cells in the follicle graph can explain and be predictive of prior COVID-19 conditions in the children.

### Position-aware SNOWFLAKE

To investigate the importance of spatial position at the single-cell level inside follicle regions, we developed another type of algorithm named position-aware SNOWFLAKE. Instead of combining morphological features after the node-level pooling layer of GNN, we integrated morphological information of both follicle and corresponding germinal center in edge features by using polar transformation from node positions (**Fig. 3a and Methods**). By adding the exact angle and distance between the cell centroid into the edge features, we restrict the graph into rigid forms that reflect closely the morphological shape of each follicle. We also add a binary feature indicating if the cell is inside or outside of the germinal center. Therefore, we have added the shape of both the follicle and germinal center in the edge and node features respectively, and provided an end-to-end predictive model.

**Fig. 3.**
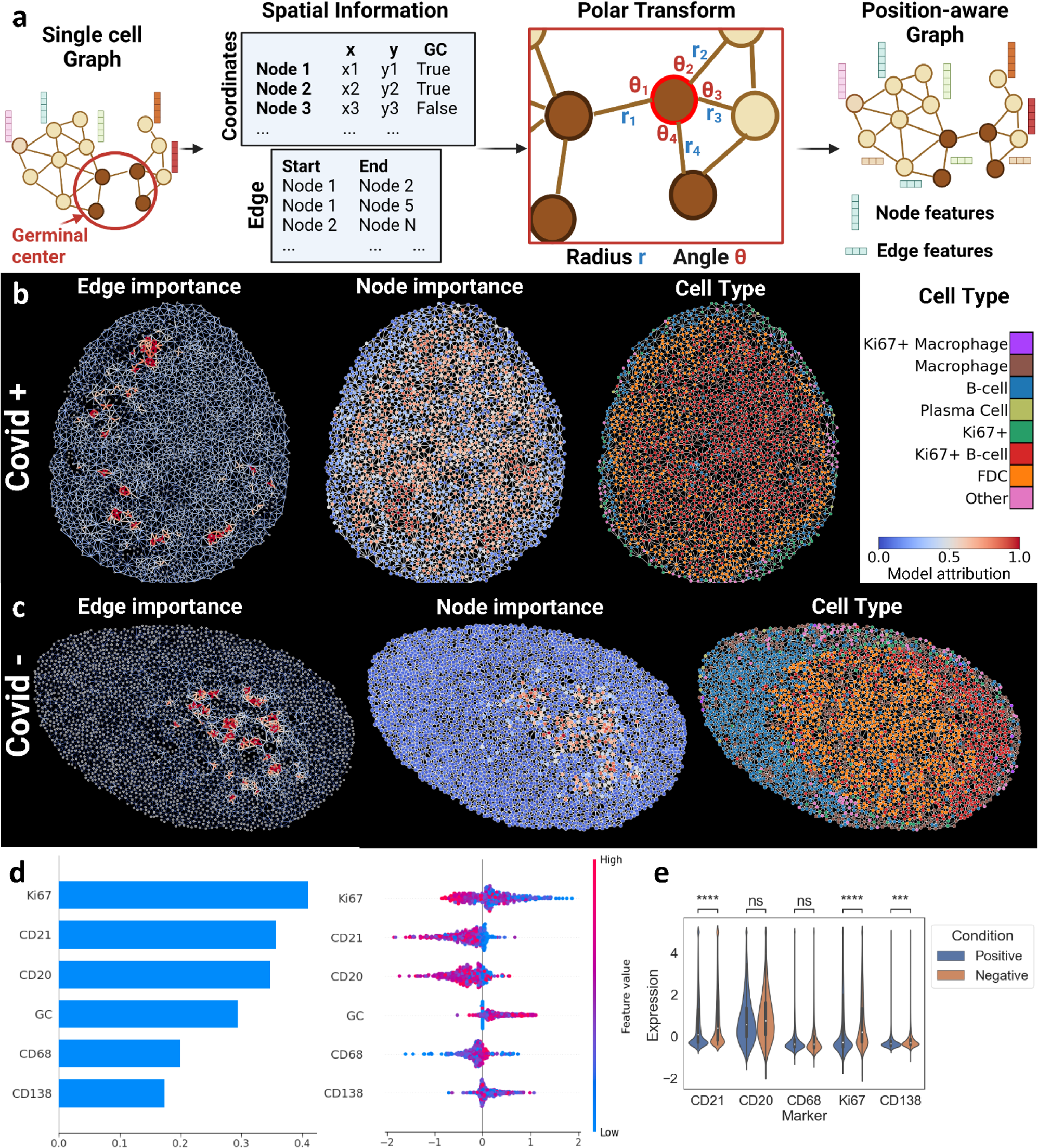
SNOWFLAKE position-aware pipeline shows important subgraphs related to patient status. **a.** Schematic showing data generation pipeline for position-aware SNOWFLAKE pipeline. Each cell’s spatial position is extracted, and the spatial graph is generated by considering the single-cell segmentation mask contact. The distance and angle of each node neighbor are calculated using polar transformation and are added as feature vectors in the edge features in the graph. **b.** Examples of edge importance (left), node importance (center), and cell type (right) projected in the original spatial domain for a COVID-positive patient’s follicle. **c.** Examples of edge importance (left), node importance (center), and cell type (right) projected in the original spatial domain for a COVID-negative patient’s follicle. **d.** Bar plot showing the importance of each node level feature for the prediction of patient-level status (left). Swarm plot showing how each node level feature expression influences SNOWFLAKE prediction (right). Each dot in the plot shows the average positive prediction or negative prediction from the node in the graphs with the colormap showing the average expression level of the nodes. **e.** Violinplot shows the comparison of single-cell marker expression distribution in follicle regions between COVID-positive and negative samples. Two-sided two-sample t-test is used for calculating p-values (ns: 0.05 < p, ***: 0.0001 < p <= 0.001, ****: p<=0.0001).

Similarly, to the previous setting, we benchmarked the position-aware SNOWFLAKE pipeline against other models. We found that position-aware SNOWFLAKE obtained similar performance as SNOWFLAKE (**Table 3**). This suggests that position-aware SNOWFLAKE can learn the morphological shape of each follicle by aggregating edge feature information. By adding the individual pairwise distance information between cells in the edge features of the graph, we are providing the same information as PCA-embedded shape embedding.

**Table 3.** SNOWFLAKE pipeline benchmarked across pooling layer and graph layer type.

| Model | Pooling Strategy | Morphology Fusion | Accuracy | AUC | F1 |
| --- | --- | --- | --- | --- | --- |
| GAT | Gated Attention | Concatenation | $0.833 \pm 0.102$ | $0.937 \pm 0.062$ | $0.875 \pm 0.064$ |
| | | Expression Only | $0.884 \pm 0.016$ | $0.953 \pm 0.013$ | $0.9 \pm 0.013$ |
| | | Position-aware | $0.856 \pm 0.077$ | **$0.96 \pm 0.021$** | $0.86 \pm 0.087$ |
| | | Tensor Decomposition | $0.842 \pm 0.078$ | $0.957 \pm 0.031$ | $0.879 \pm 0.054$ |
| | Global Attention | Concatenation | $0.821 \pm 0.081$ | $0.937 \pm 0.029$ | $0.821 \pm 0.111$ |
| | | Expression Only | $0.777 \pm 0.05$ | $0.932 \pm 0.037$ | $0.771 \pm 0.068$ |
| | | Position-aware | $0.665 \pm 0.114$ | $0.933 \pm 0.029$ | $0.576 \pm 0.194$ |
| | | Tensor Decomposition | $0.85 \pm 0.061$ | **$0.937 \pm 0.038$** | $0.882 \pm 0.042$ |
| | Max | Concatenation | $0.728 \pm 0.084$ | $0.885 \pm 0.043$ | $0.806 \pm 0.043$ |
| | | Expression Only | $0.83 \pm 0.103$ | **$0.96 \pm 0.031$** | $0.872 \pm 0.062$ |
| | | Position-aware | $0.803 \pm 0.059$ | $0.882 \pm 0.043$ | $0.822 \pm 0.053$ |
| | | Tensor Decomposition | $0.841 \pm 0.04$ | $0.878 \pm 0.053$ | $0.865 \pm 0.037$ |
| | Mean | Concatenation | $0.854 \pm 0.075$ | $0.95 \pm 0.036$ | $0.884 \pm 0.054$ |
| | | Expression Only | $0.863 \pm 0.082$ | $0.958 \pm 0.037$ | $0.892 \pm 0.059$ |
| | | Position-aware | $0.919 \pm 0.033$ | $0.972 \pm 0.019$ | $0.928 \pm 0.028$ |
| | | Tensor Decomposition | $0.926 \pm 0.01$ | **$0.973 \pm 0.009$** | $0.938 \pm 0.008$ |
| GINConv | Gated Attention | Concatenation | $0.833 \pm 0.035$ | $0.886 \pm 0.024$ | $0.858 \pm 0.032$ |
| | | Expression Only | $0.803 \pm 0.046$ | $0.885 \pm 0.016$ | $0.828 \pm 0.035$ |
| | | Position-aware | $0.853 \pm 0.017$ | **$0.911 \pm 0.007$** | $0.867 \pm 0.02$ |
| | | Tensor Decomposition | $0.802 \pm 0.04$ | $0.888 \pm 0.029$ | $0.838 \pm 0.029$ |
| | Global Attention | Concatenation | $0.803 \pm 0.019$ | $0.865 \pm 0.036$ | $0.831 \pm 0.019$ |
| | | Expression Only | $0.798 \pm 0.038$ | $0.873 \pm 0.033$ | $0.821 \pm 0.039$ |
| | | Position-aware | $0.821 \pm 0.033$ | **$0.915 \pm 0.015$** | $0.835 \pm 0.035$ |
| | | Tensor Decomposition | $0.808 \pm 0.042$ | $0.885 \pm 0.032$ | $0.824 \pm 0.047$ |
| | Max | Concatenation | $0.785 \pm 0.019$ | $0.854 \pm 0.035$ | $0.803 \pm 0.019$ |
| | | Expression Only | $0.795 \pm 0.025$ | $0.893 \pm 0.017$ | $0.82 \pm 0.028$ |
| | | Position-aware | $0.818 \pm 0.015$ | **$0.923 \pm 0.02$** | $0.842 \pm 0.01$ |
| | | Tensor Decomposition | $0.815 \pm 0.009$ | $0.875 \pm 0.024$ | $0.848 \pm 0.013$ |
| | Mean | Concatenation | $0.82 \pm 0.013$ | $0.882 \pm 0.024$ | $0.846 \pm 0.012$ |
| | | Expression Only | $0.821 \pm 0.028$ | $0.881 \pm 0.02$ | $0.845 \pm 0.026$ |
| | | Position-aware | $0.85 \pm 0.023$ | **$0.914 \pm 0.017$** | $0.867 \pm 0.022$ |
| | | Tensor Decomposition | $0.841 \pm 0.027$ | $0.892 \pm 0.024$ | $0.863 \pm 0.02$ |

More interestingly, incorporating morphological features at the edge level allows us to use attribution methods such as Integrated Gradient (IG) or GNNExplainer (Methods) on the whole graph to extract important cell subgraph biomarkers for model prediction. For each cell graph from follicles, we can extract important edges and nodes that are predictive of positive and negative COVID-19 status (**Fig. 3 b-c**). This allows us to extract subgraphs inside each follicle graph that are representative of positive and negative predictive values. Moreover, by averaging the attribution score across nodes from all graphs, we can evaluate the individual node-level features (e.g. protein marker intensity or localization) that are important predictors of prior COVID-19 infection (**Fig. 3 d**).

### SNOWFLAKE identifies recurrent spatial signatures associated with COVID-19 status

We have shown SNOWFLAKE successfully categorizes patient COVID-19 status using the combined single-cell protein expression, follicle morphological features, and corresponding single-cell neighboring graphs. Recently, multiple studies have shown the importance of both neighboring cellular composition^9,23^ as well as networks of interacting cells ^24^ in the prediction of cancer outcomes. On the other hand, a lot of attention has been given to the explainability of neural networks and more specifically graph neural networks for the understanding of node and edge importance for various tasks such as node-level prediction, link prediction, and graph classification ^16,25,26^. It is therefore critical to identify the important features and spatial single-cell subgraphs for patient-level prediction guided by our graph neural network pipeline.

SNOWFLAKE allows the identification of recurrent spatial signatures associated with COVID status from multiplex imaging by leveraging graph attribution methods (**Methods**). To understand the distribution of cell types in our graphs, we first performed unsupervised clustering from single-cell proteomic profiles to classify cell types inside follicle regions (**Supplementary Fig. 5-9 and Methods**). We compared the cell type density across follicles between COVID-positive and negative cases and found a lower density of Ki67+ germinal center B-cells in COVID-19 infected patients (**Supplementary Fig. 10a**) but with overall similar size of follicles and B-cell inside follicles showing robust GC response similar to the original paper (**Supplementary Fig. 10b**). To validate the density distribution, we compared the distribution of Ki67+ germinal center B-cells in Adenoid and Tonsil tissues and didn’t observe a statistically significant difference (**Supplementary Fig. 10c**).

To better understand and investigate subgraphs inside follicle regions that are important to COVID-19 status prediction, we developed a SNOWFLAKE explainability pipeline based on graph attribution methods. For each graph, we extracted the attribution value of nodes and edges and normalized it across the graph. Then, we extracted the subgraph induced by the high attribution nodes and edges which indicates an important subgraph with predictive values (**Methods, Fig. 4a and Supplementary Fig. 11**). For each induced subgraph, we then extracted the average attribution value and feed the subgraph to the SNOWFLAKE pipeline to obtain the subgraph COVID-19 status prediction (probability of subgraph for each condition) and extracted the subgraph embeddings from the last pooling layer of SNOWFLAKE (**Fig. 4b**).

**Fig. 4.**
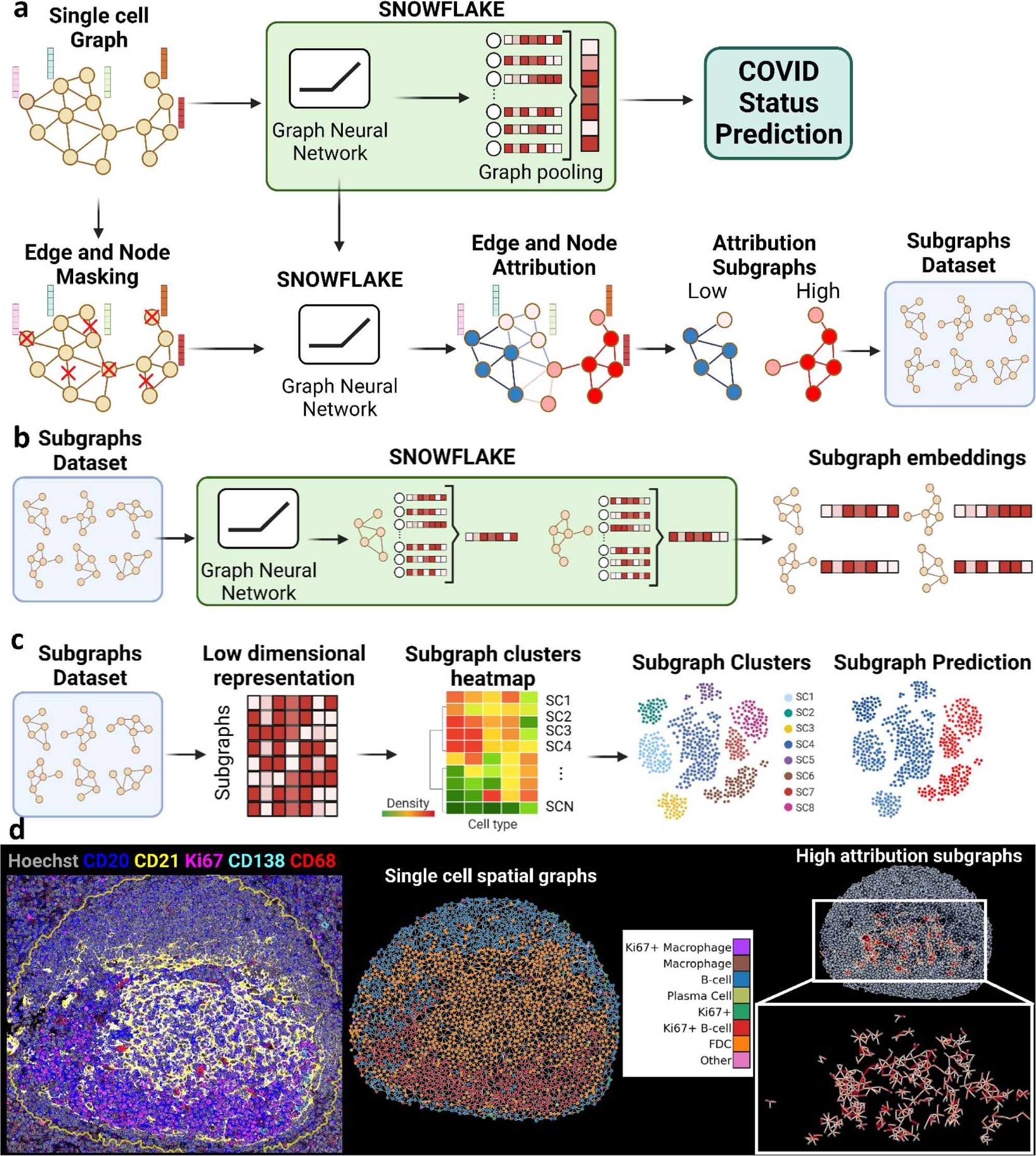
SNOWFLAKE allows extraction of subgraph biomarkers predictive of patient status. **a.** Schematic describing the extraction of high attribution subgraphs using SNOWFLAKE pipeline. First, the SNOWFLAKE model is trained on predicting COVID status from follicle imaging. Next, using the trained snowflake model, an attribution score is determined for each node and edge using masking. The corresponding high attribution edges and nodes are linked into subgraphs. Each connected component is then extracted as an individual subgraph to form a subgraph dataset. **b.** Schematic describing generation of embedding of high attribution subgraphs using SNOWFLAKE pipeline. From the subgraph dataset, each subgraph is passed through the SNOWFLAKE pipeline, and the graph embeddings of subgraphs are extracted from the last layer before the prediction head. **c.** Schematic describing subgraph clustering from low dimensional representation embedding obtained from SNOWFLAKE. The subgraphs are clustered based on their graph embedding. The subgraph clusters heatmap is visualized showing the cell type density distribution in each cluster. The corresponding subgraph predictions and clusters are visualized in a UMAP embedding. **d.** A visual example of follicle subgraph extraction pipeline. Multiplex imaging (left) of one follicle region and its contour (yellow line) is shown with Hoechst (gray), CD20 (blue), CD21 (yellow), Ki67 (magenta), CD138 (cyan), and CD68 (red) markers. The transformed single-cell spatial graph (center) is generated by transforming

Using the SNOWFLAKE embeddings of subgraphs, we can then define subgraph clusters (SC) (**Methods**) and extract the cell type density across subgraph clusters (**Fig. 4c**). This pipeline allows us to categorize various subsets of neighboring single cells into SC that consider both single cell level features and neighboring features. From multiplex imaging, the single-cell spatial graph with the corresponding cell type is generated with neighboring information, and the high attribution subgraphs can be visualized and extracted for downstream clustering at the subgraph level (**Fig. 4d**).

The corresponding subgraph cluster heatmap is visualized and each SC is associated with a probability in COVID-19 status prediction (**Fig. 5a**). Interestingly, we see a clear separation between cell types present in SC in COVID-19 positive and negative prediction (clusters 8-9-0-11-1 compared to 2-14-12-16) in certain case whereas less separation is shown in other clusters (clusters 4-10 compared to 6-5-7 and 3-15-13). Clusters 8-9-0-11-1 is a mix of B-cells with Ki67+ B-cells and macrophages. Clusters 2-14-12-16 are highly skewed towards Ki67+ B-cells and some FDCs. On the other hand, we see that clusters 3-15-13 show a mix of FDCs with other cell types such as Ki67+ B-cells, macrophage, and plasma cells, whereas clusters 4-10 and 6-5-7 both are heavily dense in FDCs but show negative and positive predictive value respectively.

**Fig. 5.**
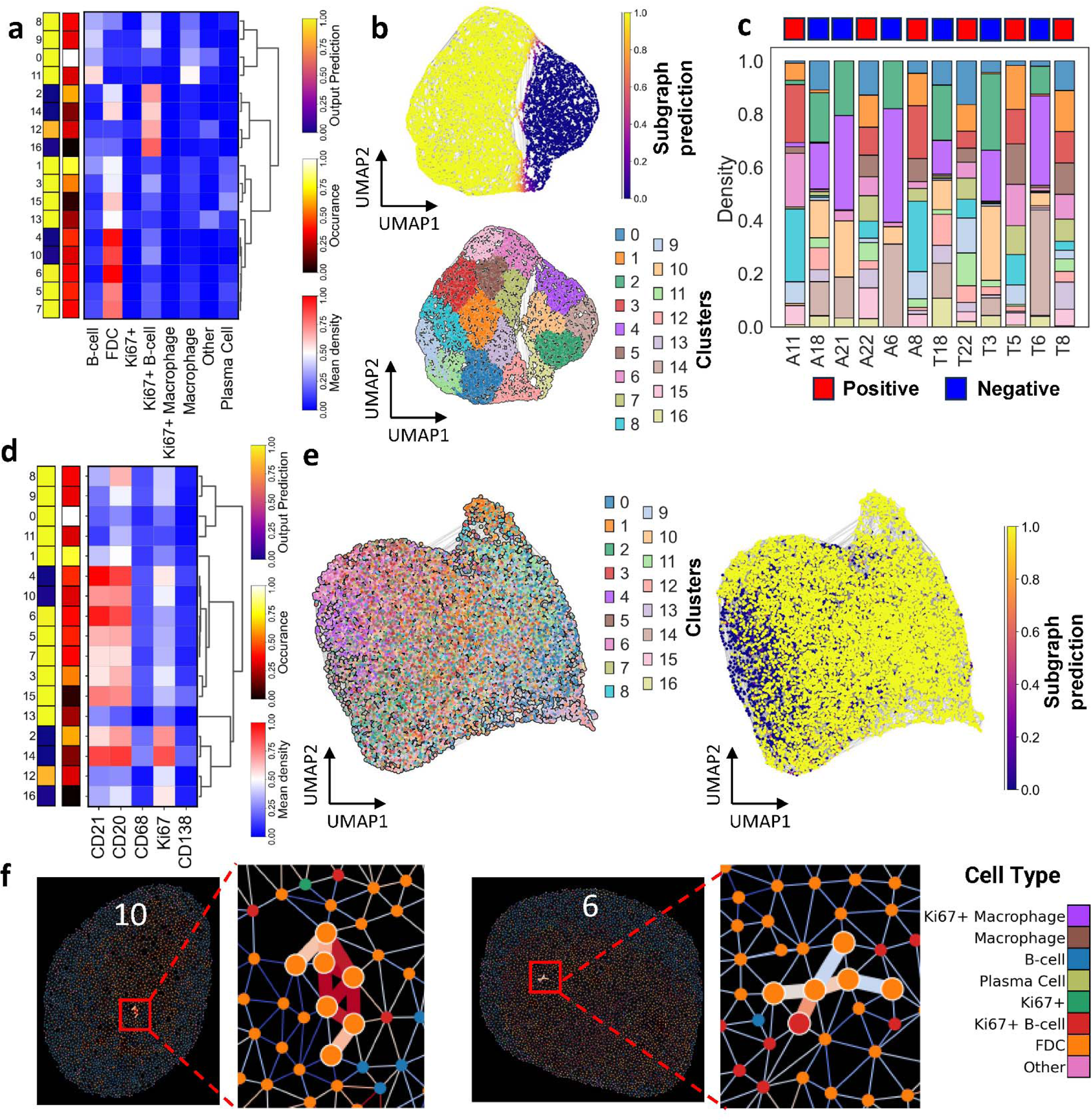
SNOWFLAKE allows extraction of subgraph biomarkers predictive of patient status. **a.** Heatmap showing the results of subgraph embedding clustering with heatmap on the right showing the relative density of each cell type in corresponding subgraph clusters (SC). The column on the left shows the subgraph-level average COVID status prediction for each SC. The column in the middle shows the relative occurrence of each SC across all the patient follicles. **b.** The two UMAPs show the SC embedding low dimensional projections with each subgraph colored by overall COVID prediction (top) and corresponding cluster (bottom). The UMAP shows a clear separation between positive and negative predictive SC. **c.** Bar plot showing the distribution of high attribution subgraph into corresponding SC of each tissue follicle region. The top colors show the patient’s COVID status (positive in red, negative in blue). **d.** Heatmap showing the results of subgraph embedding clustering with heatmap on the right showing the relative mean single-cell marker expression in corresponding subgraph clusters (SC). The column on the left shows the subgraph-level average COVID status prediction for each SC. The column in the middle shows the relative occurrence of each SC across all the patient follicles. **e.** The two UMAPs show the SC single-cell mean expression low dimensional projections with each subgraph colored by overall COVID prediction (top) and corresponding cluster (bottom). The UMAP shows a low separation between positive and negative predictive SC. **f.** Visual examples of subgraphs present in SC 10 (left) and SC 6 (right) with their corresponding edge importance and cell type in their spatial environment. The node’s color corresponds to specific cell types shown in the legend.

Furthermore, we saw a clear separation between low dimensional embedding in the UMAP space between COVID-19 positive and negative predictions (**Fig. 5b**). The SCs related to positive and negative COVID-19 status prediction are distributed across the tissue in the datasets and are not tissue-specific (**Fig. 5c**) showing the generalizability of the pipeline extracting meaningful embeddings for biomarker identification. On the other hand, when extracting the mean expression or cell type density at the subgraph level and projecting the distribution in the UMAP ^27^ low-dimension representation, we observe that SC are undifferentiable and mixed between COVID positive and negative predictions (**Fig. 5d-e, Supplementary Fig. 12 and Methods**). This shows the importance of spatial organization of single-cell in disease-status prediction and each SC represents a unique set of single-cell organization extracted from the SNOWFLAKE pipeline (**Supplementary Fig. 13**).

We looked at the spatial connectivity inside high and low-prediction SCs to understand the spatial motifs important for predicting cell information (**Fig. 6a**). For each SC, we looked at the pairwise connection density between cell types by counting the number of edges between cell type and then we aggregate the connectivity information across SC in a heatmap format (**Fig. 6b**). When comparing the connection present in SC 2, SC 6, and SC 10, we can see that the SC 6 and 10 both contain a high number of FDCs with SC 10 having more Ki67+ B-cells connection with whereas SC 6 is more have some macrophages connectivity too. SC2 is more uniformly distributed between Ki67+ B-cells and FDCs connections (**Fig. 5c**). Similarly, we plotted the connectivity graphs across similar clusters to understand the different motifs of cell type more recurrent in COVID-positive and negative cases (**Supplementary Fig. 14**).

**Fig. 6.**
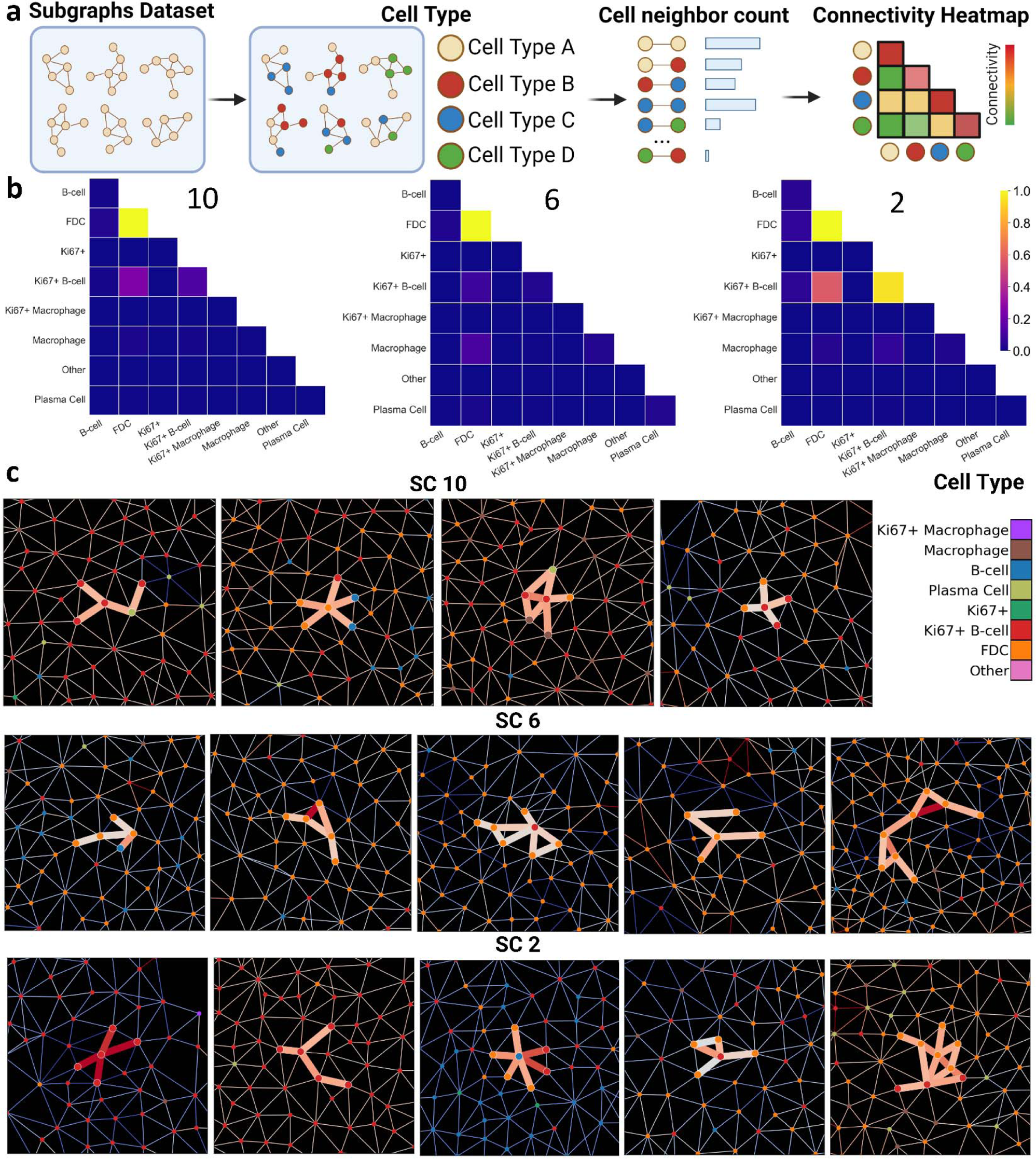
SNOWFLAKE allows extraction of subgraph biomarkers predictive of patient status. **a.** Schematic showing the subgraph cluster connectivity analysis. From each subgraph cluster, all the subgraph nodes corresponding cell types are extracted as well as their neighboring information. The pairwise neighboring cell type connection frequency is calculated by counting the number of edges between the corresponding cell types. All this information is then represented as a connectivity heatmap showing the relative connectivity strength between specific cell types. **b.** Edge connectivity distribution of SC 10 (left), SC 6 (center), and SC 2 (right) showing which cell type interacts with each other mode often in each SC. **c.** Visual example showing the distribution of subgraphs present in SC 10 (top), SC 6 (left), and SC2 (bottom) with their corresponding edge importance and cell type in their spatial environment. The node’s color corresponds to specific cell types shown in the legend. The edge color shows the attribution score of each edge extracted from the SNOWFLAKE pipeline.

## Discussion

In this study, we developed SNOWFLAKE, a multi-modal graph-based learning method combining spatial morpho-proteomic features (both follicle morphology and multiplexed proteomic markers of cell compositions) in a single pipeline within multiplexed proteomic images of lymphoid tissues. Leveraging the single-cell spatial organization and multiplex protein markers, SNOWFLAKE successfully predicted the patient-level COVID-19 exposure from follicle-unique features. We benchmarked the importance of morphological and spatial features in our predictive pipeline and showed the predictive superiority of the SNOWFLAKE pipeline.

To better understand the important cellular motifs from the SNOWFLAKE pipeline, we utilized attribution and explanation methods for extracting highly important sets of edges and nodes from our cellular graphs in the follicle regions. We further clustered the resulting high-importance subgraphs into distinct subgraph clusters using graph embeddings from the SNOWFLAKE pipeline to classify the subgraph clusters into disease-relevant predictive biomarkers. We showed that the SNOWFLAKE pipeline successfully extracts embeddings from graph organization for stratification of subgraphs into positive and negative predictive motifs which are not possible when only considering subgraph marker expression or cell type density. More importantly, the extracted cell motifs are inferred from the SNOWFLAKE pipeline which allows diverse size and connection patterns to be considered (**Fig. 7**). This has biological importance in understanding the longer-standing effects of COVID-19 in the convalescent phase.

**Fig. 7.**
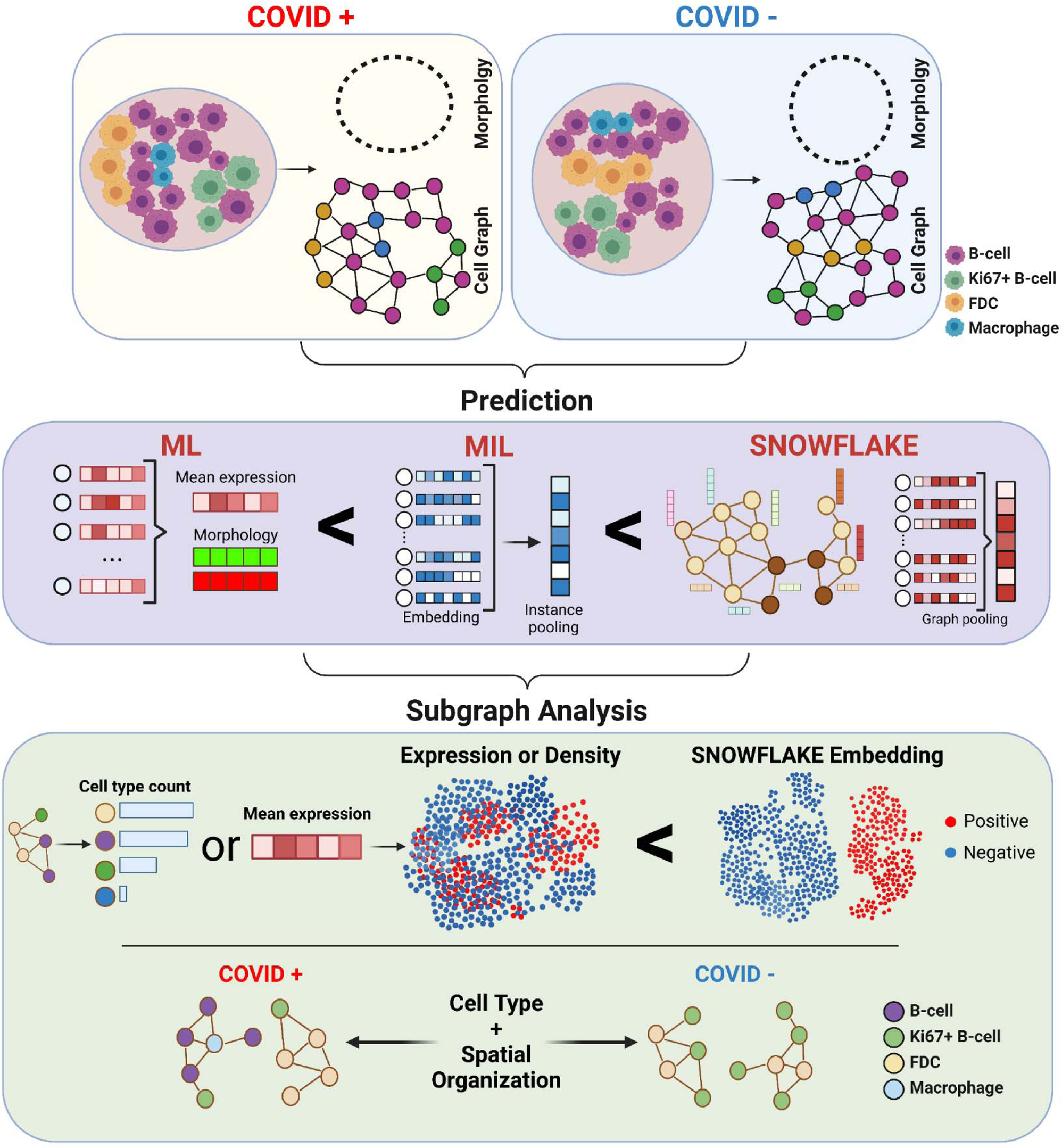
SNOWFLAKE enables higher disease status prediction accuracy with spatial biomarker explainability. SNOWFLAKE pipeline was used to classify B-cell follicle structures from COVID-positive and negative patients. We have shown better predictive abilities of our model when combining morphological and spatial information of single-cell microenvironment structures compared to expression only. Moreover, using trained graph neural network and attribution methods, we extracted high predictive value subgraphs and compared the spatial connectivity between subgraph clusters with COVID positive and negative predictive values.

We showed in this work the importance of modeling single-cell multiplex data as cell neighborhood graphs to predict patient-level information. Single-cell level neighborhood connectivity is indicative of patient disease status. One limitation of this study is the limited number of protein markers for the identification of cell types inside our follicle database. It is also hard to combine different imaging databases and imaging modalities due to the difference between multiplex marker panels and imaging technologies. Novel computation methods for unseen spatial single-cell dataset annotation ^28,29^ and missing marker imputation ^30,31^ could be used to further improve the generalizability of the SNOWFLAKE pipeline. Despite these limitations, SNOWFLAKE provides a complementary solution to the need to extract relevant single-cell motifs prediction and biomarker identification for the understanding of spatial arrangement in health and disease. SNOWFLAKE is a generalizable pipeline for understanding disease-related features from single-cell multiplex spatial omics data and is applicable to various studies such as vaccine response, malignant lymph nodes, tertiary lymphoid structures, or other type of diseases such as cancer and glomeruli in normal and diseased kidneys.

SNOWFLAKE is a predictive pipeline that incorporates morphological and marker expression features from single-cell level imaging data. Deep learning guided cell local graph identification is used to characterize spatial motifs that are related to health and disease. The corresponding local graphs are further stratified using their embeddings to highlight the relationship between motif classes and patient-level status. With the advancement of multiplex imaging throughput for spatial proteomics and transcriptomics, SNOWFLAKE allows the investigation of important cellular neighboring graphs with patient status.

## Methods

### Curation of the dataset for SNOWFLAKE

We created a database of human B-cell follicles from various lymphoid tissues from various multiplex protein datasets from both public data repositories and published papers. Multiplex technologies were used including Cyclic Immunofluorescence (CycIF) ^32–34^, CO-Detection by indEXing (CODEX) ^17,35^ for a total of 930 follicles and 775 germinal centers **(Supplementary Table 1).** The primary follicles and the germinal centers in these datasets were manually segmented using DAPI, CD20, CD21, and Ki67 signals. In the NIH-COVID dataset, we have a total of 442 follicles with 391 germinal centers with 253 follicles (and 219 germinal centers) from COVID-positive patients and 189 (172 germinal centers) from COVID-negative patients (**Table 1**). Sample collection was approved by the Institutional Review Board (IRB) at Children’s National Hospital (IRB protocol number 00009806).

### Single-cell segmentation

To obtain an accurate segmentation in follicle regions with dense cells, we performed a two-step semi-supervised segmentation approach. First, we manually labeled pixel data using iLastik (v 1.4.0) to predict the pixels corresponding to the nucleus, cytoplasmic, and background using the combination of protein channel images. A probability map of three labels was created corresponding to the nucleus, cytosol, and background for each tissue is generated with manual annotation of a few regions for each label. These probability maps were then added to the CellProfiler (v 4.2.1) software for the single-cell segmentation. This step is critical because it generates the single-cell masks that were later used for all the downstream analysis. The pipeline starts with identifying the primary objecting or the nuclei using the “IdentifyPrimaryObjects” module and setting the diameter limits to 8-13 pixels. Then, the cellular membranes were identified using the “IdentifySecondaryObjects” module by expanding the primary objects along the nuclear+cytoplasm probabilities. Finally, the cellular membranes were used to generate the single-cell masks that were later exported as tiff images. Single cells were segmented from the whole tissue area and assigned to the follicle based on the location of cell centroids inside or outside a specific follicle.

### Single-cell unsupervised clustering

Single-cell unsupervised clustering was performed using Leiden algorithm ^36^ a graph-based community detection algorithm from all tissue images. From each segmented cell region, the mean intensity of each marker expression was calculated. The resulting feature matrix consisted of n rows of the total number of cells and p columns of marker expression. Each column of the feature matrix was normalized z-score and batch correction between samples was performed using Scanorama pipeline ^37^. The neighborhood graph in the embedding space was constructed and used for unsupervised community detection. After the first round of unsupervised clustering, we annotated each cluster (n_cluster = 24), and we merged similar clusters to obtain 11 final clusters. For analysis, we extracted cells inside follicle regions only which resulted in cell type distribution in 10 clusters.

### Follicle and germinal center segmentation

Follicle and germinal center regions were manually segmented by looking at multiplex protein markers such as CD20, CD21, Ki67, and Hoechst. CD20 and Hoechst were mainly used to determine the follicle regions in lymphoid tissues. While CD20 staining allows the identification of the B-cell membrane, the Hoechst marker shows good separation between the mantle zone (compact zone) and the germinal center (darker zone) in follicle regions. Ki67 and CD21 were used for the characterization of high proliferation cells and follicular dendritic cells respectively for identification of germinal center regions. Therefore, we obtained an accurate segmentation of follicles and corresponding germinal centers in the tissues. **Follicle and germinal center morphology analysis:** After manual segmentation of follicles and germinal center regions, we extract boundaries of follicles and germinal centers by taking the contour of region masks using OpenCV findContours function transforming manual segmentation regions into a set of points defining the contour of follicles and germinal center regions. Next, we performed Procrustes analysis for shape alignment, scaling, and shift ^18^, we resampled the contour points into a vector of equally spaced points (*n_points_* = 120) using B-spline representation along each contour. We processed each follicle and corresponding germinal center by pair since their morphology is related. The sampled coordinates for each follicle and corresponding germinal center are normalized to the center location (0,0) by subtracting the mean coordinate of the follicles. Since follicles have different sizes and diameters, we decoupled the shape and size of the follicles by dividing the coordinates of follicles and germinal center by the scale of shape 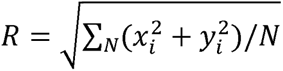 of each follicle. Each follicle and corresponding germinal center coordinates are rotated along the major axis of the follicle coordinates using Singular Value Decomposition. Finally, a fine-tuned rotation minimization step is performed for each follicle and average follicle shape across the dataset ^38^. Principal component analysis (PCA) was then applied to these descriptors for all follicles to obtain eigenshape vectors. The first 16 principal components from the eigenshape vectors were used as a simplification set of descriptors for follicle and germinal center shapes. This was determined by the reconstruction shape and variance explained. We benchmarked MorphPCA against an Auto Encoder (AE) type of deep learning architecture for morphology embedding of follicles and germinal centers. We used Hausdorff distance and Mean Squarred Error to quantify the reconstruction of follicles and germinal center morphology from low dimensional embeddings (**Supplementary Fig. 15**).

### Single-cell graph construction

We extract the corresponding 2D localization for each segmented cell centroid and assign a node in the created graph. The node features are assigned by creating a vector representing the mean protein expression value per cell. Cell spatial graphs are created using cell contact information. Cell contacts are calculated by binary dilatation of single-cell masks and cells that overlap were considered to have contact ^39^. We choose cell mask contact for spatial neighboring maps generation in tonsil samples because of the higher density of cells. The models are trained in a multi-instance learning framework, that is a follicle label for each instance is assigned based on the condition of the tissue.

### Graph neural networks

For the graph neural network, we used a 2-layer network of embedding size 16. The input of the model is the generated single-cell graph for each follicle with node feature represented by a vector of marker mean intensities. The input of each node is first passed through a dense layer to learn embedding representation. We benchmarked different graph layers’ backbones (GCN^40^, GAT^41^, GINConv^42^, GraphConv^43^, and SAGEConv^44^) for optimal prediction in the validation set (**Supplementary Table 3**). Moreover, we use LayerNorm followed by activation for each layer to prevent overfitting and use a rectified linear unit (PReLU) as the activation layer and residual connection between each layer to improve the learning capabilities and stability ^45,46^. Finally, the node embeddings are then aggregated by a pooling layer and two dense layers were then used to obtain prediction at the follicle level (i.e., graph-level prediction).

### Multi-instance learning baseline

A MIL multi-layer perception baseline was used to compare our graph neural network. In the same multi-instance learning framework, we assign a class label from the follicle condition. The input of the model is the generated single-cell graph for each follicle with node feature represented by a vector of marker mean intensities. Here, we use stacked dense layers of embedding size 16 to obtain a node embedding. Each layer transforms the input as the following function: *H*^*l*+1^ = *f*(*W^l^H^l^*) with *l* the corresponding layer, *f* the activation function, *H^l^* the node embedding matrix and *W^l^* the weight matrix of the layer *l*. Finally, the node embeddings are then aggregated by a pooling layer and two dense layers were then used to obtain prediction at the follicle level (i.e., graph-level prediction).

### SNOWFLAKE pipeline

SNOWFLAKE pipeline consists of the integration of single-cell level features with follicle-level morphological features for the follicle level prediction (COVID status). For this purpose, we aggregated graph or MIL node-level features using a pooling layer and combined expression features and morphological features using modality fusion. We used a pooling layer to aggregate the node-level embedding for graph-level prediction. We benchmarked various pooling layers to investigate the best way of incorporating node-level features: mean, max, sum, global attention ^47^, and gated attention ^48^ (**Supplementary Table 3**). After the graph pooling layer, we obtain a graph-level embedding used for prediction. To incorporate morphological features, we used modality fusion using a tensor decomposition fusion module via Kronecker Product ^49^ with a gating-based attention mechanism ^50^. Modality fusion is benchmarked against the concatenation of graph and morphological features.

### SNOWFLAKE position-aware

To compare the effect of modality integration before and after the pooling layer of our GNN, we propose a position-aware SNOWFLAKE pipeline. Instead of integrating morphology information from PCA analysis, we directly include the polar transformation based on each node’s spatial location in the graph edge weights. That is, from each pair of connected nodes in the spatial graph created from single cells, we extracted ρ,θ which represents the distance and angular rotation between the two nodes respectively. For each edge in our graph, we obtain a vector of size two representing the distance and rotation between the connected nodes. The corresponding vector is used to define the features of each edge.

### Multi-layer perception and Machine Learning baseline

For benchmarking SNOWFLAKE, we trained an MLP model on the follicle mean expression of all protein markers using the same embedding dimension and number of layers. On the other hand, several machine learning models were used for a baseline on mean follicle expression level which includes Naïve Bayes, Random Forest, AdaBoost, Decision Tree, Support Vector Machine (SVM), and Gradient Boosting. We used the scikit-learn python library with default setting when training and testing these machine learning models. We have also tested prediction using morphology features only and mean expression combined with morphology features. In total, we have benchmarked prediction from mean expression only (1), morphology features only (2), and combined mean expression with morphology features (3) per follicle.

### Feature interpretability and subgraph extraction

To characterize the feature importance magnitude and direction of impact we used SHAP ^51^, Integrated Gradient (IG) ^15^, and GNNExplainer ^16^. SHAP, IG, and GNNExplainer provide local interpretations as attribution scores of how each feature in each cell or each node/edge (in graphs) contributes to the single instance (follicle-level) model prediction. The mean absolute value of the attribution scores shows the global impact on model prediction. The individual node (cell) or edge level attribution scores can also be used to rank the importance of each corresponding node and edge toward the model prediction but also understand the contribution for prediction being positive or negative. From the SNOWFLAKE pipeline, node and edge level attribution scores are calculated with a subset of nodes and edges extracted based on a threshold (for example top or bottom 10%). The corresponding induced subgraphs are obtained from the extracted nodes and edges.

### Subgraph clustering and connectivity analysis

The subgraphs extracted from attribution scores are clusters to understand the corresponding SNOWFLAKE-induced single-cell motifs. Each subgraph embedding is calculated using the trained SNOWFLAKE pipeline’s last layer before the prediction head. This provides us with low-dimensional embeddings of the same size induced by model prediction for graphs with various numbers of nodes and edges. The low embeddings are then used to cluster subgraphs into subgraph clusters (SC) and understand the overall predictive ability of each SC. We compared the separation between the positive and negative predictive subgraph using embedding from SNOWFLAKE embedding, mean cell marker expression level, and mean cell type densities.

### Prediction metrics

To compare the models’ prediction abilities, we used a 5-fold cross-validation setting by separating the dataset into an 80% training set and a 20% validation set. We used Accuracy, ROC AUC score, and F1 score as metrics to evaluate the data prediction in the validation sets. When characterizing our model performance, we focused on the ROC AUC score which is more suited for class imbalance problems.

### Statistical testing

The details of statistical tests employed in each case were provided in the figure captions. All P values were corrected for multiple testing and the statistical testing method was indicated in the figure captions. We used the following convention to indicate significance with asterisks: not significant (ns) (P > 0.1), * (0.1 > P > 0.01), ** (0.01 > P > 0.001), *** (0.001 > P > 0.0001), and **** (P ≤ 0.0001).

## Acknowledgments

A.F.C. holds a career award at the Scientific Interface from Burroughs Wellcome Fund and a Bernie-Marcus Early-Career Professorship. A.F.C. was supported by start-up funds from the Georgia Institute of Technology and Emory University. Research reported in this study was supported by the National Institute of General Medical Sciences of the National Institutes of Health under award number R35GM151028 and 1R21AI173900. This work was supported in part by intramural funding from the National Institute of Allergy and Infectious Diseases (NIAID), NIH and Andrea J. Radtke, Ronald N. Germain from Center of Advance Tissue Imaging.

## Authors contributions

TH contributed to the data processing, data analysis, and writing of this paper. MA contributed to the writing of this paper, VK contributed to data processing. SLG, QX, PM, and KM contributed to the data. AFC supervised the project and wrote the paper.

## Data and materials availability

All data needed to evaluate the conclusions in the paper are present in the paper and/or the Supplementary Materials. The SNOWFLAKE source code, the training and test data, as well as the trained SNOWFLAKE models, are available upon request.

## References

1. Nichols, H. B. Snow Crystals: Natural and Artificial. Ukichiro Nakaya. Harvard Univ. Press, Cambridge, 1954. xii + 510 pp. Illus. $10. Science 120, 755–755 (1954).

2. Demange, G., Zapolsky, H., Patte, R. & Brunel, M. A phase field model for snow crystal growth in three dimensions. Npj Comput. Mater. 3, 15 (2017).

3. Pereira, J. P., Kelly, L. M. & Cyster, J. G. Finding the right niche: B-cell migration in the early phases of T-dependent antibody responses. Int. Immunol. 22, 413–419 (2010).

4. Stebegg, M. et al. Regulation of the Germinal Center Response. Front. Immunol. 9, 2469 (2018).

5. Elsner, R. A. & Shlomchik, M. J. Germinal Center and Extrafollicular B Cell Responses in Vaccination, Immunity, and Autoimmunity. Immunity 53, 1136–1150 (2020).

6. Allen, C. D. C. & Cyster, J. G. Follicular dendritic cell networks of primary follicles and germinal centers: Phenotype and function. Semin. Immunol. 20, 14–25 (2008).

7. Wang, X. et al. Follicular dendritic cells help establish follicle identity and promote B cell retention in germinal centers. J. Exp. Med. 208, 2497–2510 (2011).

8. Zhou, J., et al. Graph Neural Networks: A Review of Methods and Applications. ArXiv181208434 Cs Stat (2021).

9. Wu, Z. et al. Graph deep learning for the characterization of tumour microenvironments from spatial protein profiles in tissue specimens. Nat. Biomed. Eng. 6, 1435–1448 (2022).

10. Wang, Y. et al. Cell graph neural networks enable the precise prediction of patient survival in gastric cancer. Npj Precis. Oncol. 6, 45 (2022).

11. Pati, P. et al. Hierarchical graph representations in digital pathology. Med. Image Anal. 75, 102264 (2022).

12. Nair, A. et al. A graph neural network framework for mapping histological topology in oral mucosal tissue. BMC Bioinformatics 23, 506 (2022).

13. Govek, K. W. et al. CAJAL enables analysis and integration of single-cell morphological data using metric geometry. Nat. Commun. 14, 3672 (2023).

14. Li, Y., Nowak, C. M., Pham, U., Nguyen, K. & Bleris, L. Cell morphology-based machine learning models for human cell state classification. Npj Syst. Biol. Appl. 7, 23 (2021).

15. Sundararajan, M., Taly, A. & Yan, Q. Axiomatic Attribution for Deep Networks. ArXiv170301365 Cs (2017).

16. Ying, R., Bourgeois, D., You, J., Zitnik, M. & Leskovec, J. GNNExplainer: Generating Explanations for Graph Neural Networks. Preprint at http://arxiv.org/abs/1903.03894 (2019).

17. Xu, Q. et al. Adaptive immune responses to SARS-CoV-2 persist in the pharyngeal lymphoid tissue of children. Nat. Immunol. 24, 186–199 (2023).

18. Phillip, J. M., Han, K.-S., Chen, W.-C., Wirtz, D. & Wu, P.-H. A robust unsupervised machine-learning method to quantify the morphological heterogeneity of cells and nuclei. Nat. Protoc. 16, 754–774 (2021).

19. Sokal, A. et al. Maturation and persistence of the anti-SARS-CoV-2 memory B cell response. Cell 184, 1201–1213.e14 (2021).

20. Gaebler, C. et al. Evolution of antibody immunity to SARS-CoV-2. Nature 591, 639–644 (2021).

21. Chertow, D., et al. SARS-CoV-2 infection and persistence throughout the human body and brain. https://www.researchsquare.com/article/rs-1139035/v1 (2021) doi:10.21203/rs.3.rs-1139035/v1.

22. Laidlaw, B. J. & Ellebedy, A. H. The germinal centre B cell response to SARS-CoV-2. Nat. Rev. Immunol. 22, 7–18 (2022).

23. Sorin, M. et al. Single-cell spatial landscapes of the lung tumour immune microenvironment. Nature 614, 548–554 (2023).

24. Gaglia, G. et al. Lymphocyte networks are dynamic cellular communities in the immunoregulatory landscape of lung adenocarcinoma. Cancer Cell 41, 871–886.e10 (2023).

25. Agarwal, C., Queen, O., Lakkaraju, H. & Zitnik, M. Evaluating Explainability for Graph Neural Networks. Preprint at http://arxiv.org/abs/2208.09339 (2023).

26. Rathee, M., Funke, T., Anand, A. & Khosla, M. BAGEL: A Benchmark for Assessing Graph Neural Network Explanations. Preprint at http://arxiv.org/abs/2206.13983 (2022).

27. Becht, E. et al. Dimensionality reduction for visualizing single-cell data using UMAP. Nat. Biotechnol. 37, 38–44 (2019).

28. Brbić, M. et al. Annotation of spatially resolved single-cell data with STELLAR. Nat. Methods 19, 1411–1418 (2022).

29. Shaban, M., et al. MAPS: Pathologist-level cell type annotation from tissue images through machine learning. http://biorxiv.org/lookup/doi/10.1101/2023.06.25.546474(2023) doi:10.1101/2023.06.25.546474.

30. Wu, E. et al. 7-UP: Generating in silico CODEX from a small set of immunofluorescence markers. PNAS Nexus 2, pgad171 (2023).

31. Ghahremani, P., et al. DeepLIIF: Deep Learning-Inferred Multiplex ImmunoFluorescence for IHC Image Quantification. http://biorxiv.org/lookup/doi/10.1101/2021.05.01.442219 (2021) doi:10.1101/2021.05.01.442219.

32. Rashid, R. et al. Highly multiplexed immunofluorescence images and single-cell data of immune markers in tonsil and lung cancer. Sci. Data 6, 323 (2019).

33. Schapiro, D. et al. MCMICRO: a scalable, modular image-processing pipeline for multiplexed tissue imaging. Nat. Methods 19, 311–315 (2022).

34. Lin, J.-R. et al. Highly multiplexed immunofluorescence imaging of human tissues and tumors using t-CyCIF and conventional optical microscopes. eLife 7, e31657 (2018).

35. Black, S. et al. CODEX multiplexed tissue imaging with DNA-conjugated antibodies. Nat. Protoc. 16, 3802–3835 (2021).

36. Traag, V. A., Waltman, L. & van Eck, N. J. From Louvain to Leiden: guaranteeing well-connected communities. Sci. Rep. 9, 5233 (2019).

37. Hie, B., Bryson, B. & Berger, B. Efficient integration of heterogeneous single-cell transcriptomes using Scanorama. Nat. Biotechnol. 37, 685–691 (2019).

38. Wu, P.-H. et al. Evolution of cellular morpho-phenotypes in cancer metastasis. Sci. Rep. 5, 18437 (2016).

39. Martinelli, A. L. & Rapsomaniki, M. A. ATHENA: analysis of tumor heterogeneity from spatial omics measurements. Bioinformatics 38, 3151–3153 (2022).

40. Kipf, T. N. & Welling, M. Semi-Supervised Classification with Graph Convolutional Networks. Preprint at http://arxiv.org/abs/1609.02907 (2017).

41. Veličković, P. et al. Graph Attention Networks. Preprint at http://arxiv.org/abs/1710.10903 (2018).

42. Xu, K., Hu, W., Leskovec, J. & Jegelka, S. How Powerful are Graph Neural Networks? Preprint at http://arxiv.org/abs/1810.00826 (2019).

43. Morris, C. et al. Weisfeiler and Leman Go Neural: Higher-order Graph Neural Networks. Preprint at http://arxiv.org/abs/1810.02244 (2021).

44. Hamilton, W. L., Ying, R. & Leskovec, J. Inductive Representation Learning on Large Graphs. ArXiv170602216 Cs Stat (2018).

45. You, J., Ying, R. & Leskovec, J. Design Space for Graph Neural Networks. Preprint at http://arxiv.org/abs/2011.08843 (2021).

46. Ioffe, S. & Szegedy, C. Batch Normalization: Accelerating Deep Network Training by Reducing Internal Covariate Shift. Preprint at http://arxiv.org/abs/1502.03167 (2015).

47. Li, Y., Tarlow, D., Brockschmidt, M. & Zemel, R. Gated Graph Sequence Neural Networks. (2015) doi:10.48550/ARXIV.1511.05493.

48. Ilse, M., Tomczak, J. M. & Welling, M. Attention-based Deep Multiple Instance Learning. ArXiv180204712 Cs Stat (2018).

49. Hu, G. et al. Attribute-Enhanced Face Recognition with Neural Tensor Fusion Networks. in 2017 IEEE International Conference on Computer Vision (ICCV) 3764–3773 (IEEE, 2017). doi:10.1109/ICCV.2017.404.

50. Chen, R. J. et al. Pathomic Fusion: An Integrated Framework for Fusing Histopathology and Genomic Features for Cancer Diagnosis and Prognosis. ArXiv191208937 Cs Q-Bio (2020).

51. Lundberg, S. M. & Lee, S.-I. A Unified Approach to Interpreting Model Predictions. in Advances in Neural Information Processing Systems (eds. Guyon, I. et al.) vol. 30 (Curran Associates, Inc., 2017).

