## Supplementary Information for "Spatial Morphoproteomic Features Predict Uniqueness of Immune Microarchitectures and Responses in Lymphoid Follicles"

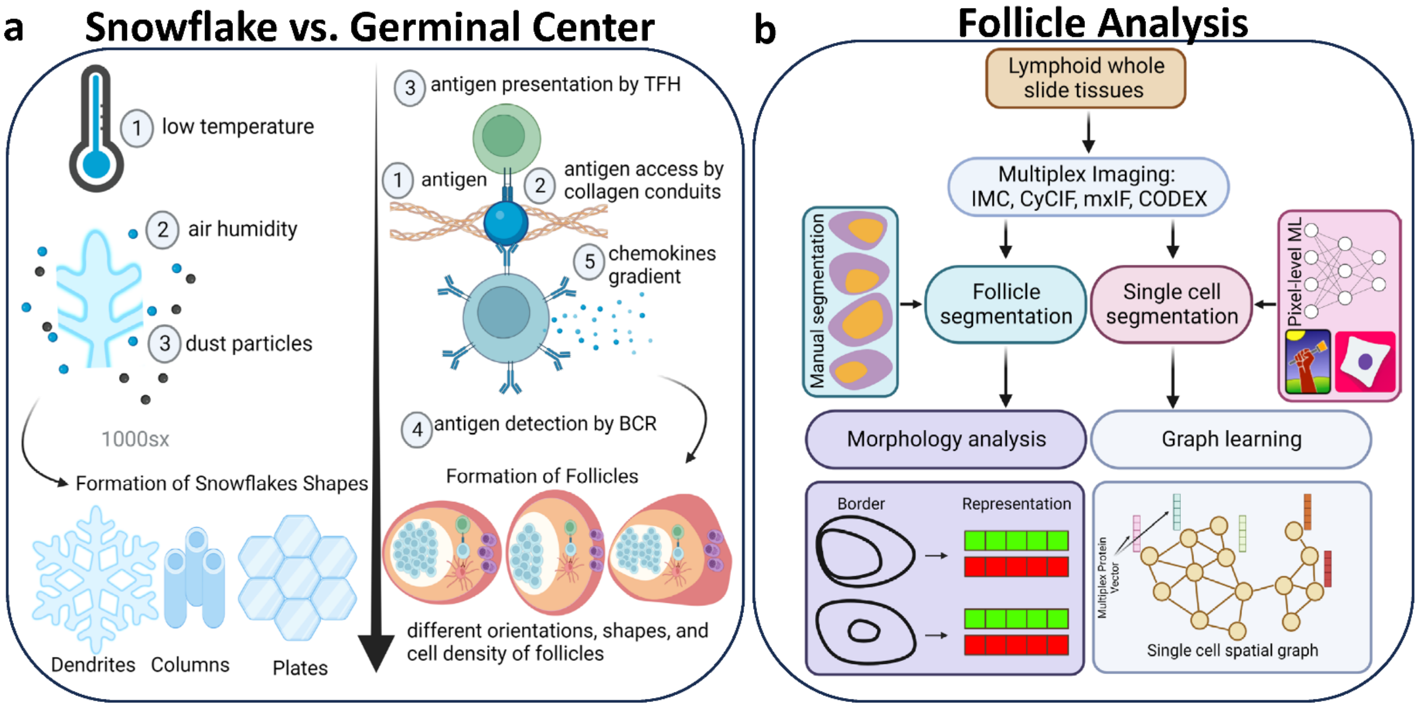
a
Snowflake vs. Germinal Center
1 low temperature
2 air humidity
3 dust particles
1000sx
Formation of Snowflakes Shapes
Dendrites
Columns
Plates
3 antigen presentation by TFH
1 antigen
2 antigen access by collagen conduits
5 chemokines gradient
4 antigen detection by BCR
Formation of Follicles
different orientations, shapes, and cell density of follicles
b
Follicle Analysis
Lymphoid whole slide tissues
Multiplex Imaging: IMC, CyCIF, mxIF, CODEX
Manual segmentation
Follicle segmentation
Single cell segmentation
Pixel-level ML
Morphology analysis
Graph learning
Border
Representation
Multiplex Protein Vector
Single cell spatial graph

**Supplementary Fig. 1. Overview of the processing pipeline for SNOWFLAKE.**

**a.** Comparison of parallel between snowflake formation with B-cell follicle germinal center formation.

**b.** Overview of Follicle analysis pipeline. From lymphoid whole slide tissues, multiplex imaging technologies are used to image targeted protein biomarkers. Follicle segmentation and single-cell segmentation are performed in parallel which produces morphological analysis for follicle segmentations and graph learning analysis from single-cell segmentations.

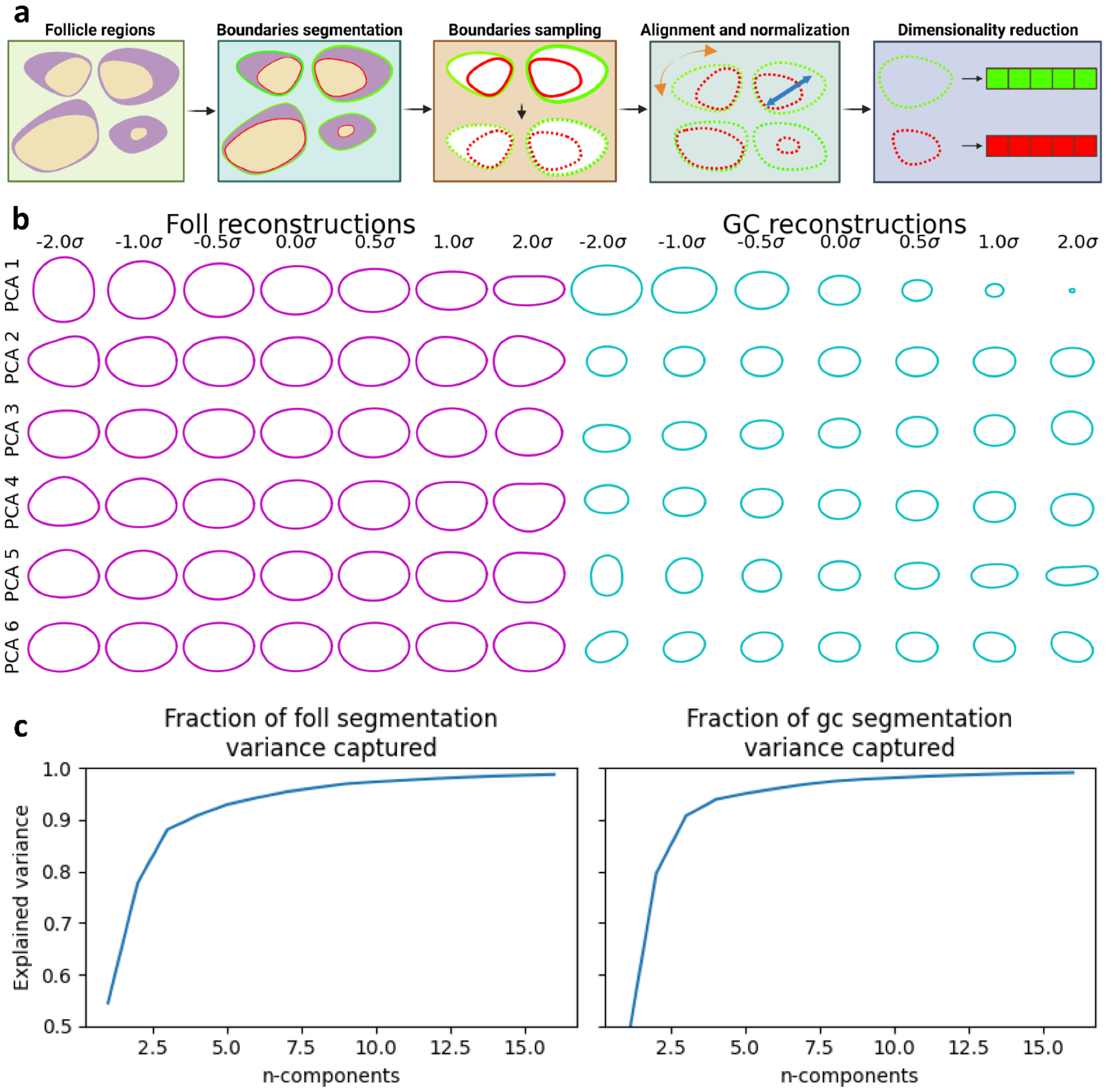
a
Follicle regions
Boundaries segmentation
Boundaries sampling
Alignment and normalization
Dimensionality reduction
b
Foll reconstructions
GC reconstructions
-2.0σ
-1.0σ
-0.5σ
0.0σ
0.5σ
1.0σ
2.0σ
PCA 1
PCA 2
PCA 3
PCA 4
PCA 5
PCA 6
c
Fraction of foll segmentation variance captured
Fraction of gc segmentation variance captured
Explained variance
n-components
1.0
0.9
0.8
0.7
0.6
0.5
2.5
5.0
7.5
10.0
12.5
15.0

**Supplementary Fig. 2. Overview of morphology analysis of follicle and germinal centers.**

**a.** Schematic showing the morphology analysis pipeline of follicle regions. Boundaries of follicles and germinal centers are segmented based on multiplex imaging. An equal number of coordinate points (N=160) are sampled along the boundaries equally spaced for follicle and germinal center boundaries. The sampled coordinates are aligned with respect to their major axis and normalized based on the scale of shape value. PCA transformation is used to reduce the aligned and normalized coordinate vectors into a set of principal components (n=16) capturing most of the variation of the follicle and germinal center shapes

**b.** Plot showing the variation of the follicle (left) and germinal center (right) shapes across their respective 6 first principal components. Each row shows the variation of the shape along one principal component. Each column represents a multiple (-2, -1, -0.5, 0, 0.5, 1, 2) of the standard deviation along one principal component. For each case, only one principal component value is changed while all other principal component values are fixed to be equal to their respective means.

**c.** Line plot showing the variation of explained variance as a function of the number of PCA components for follicle segmentation (left) and germinal center segmentation (right). The Y-axis shows the fraction of variance explained and the X-axis shows the number of principal components from PCA considered.

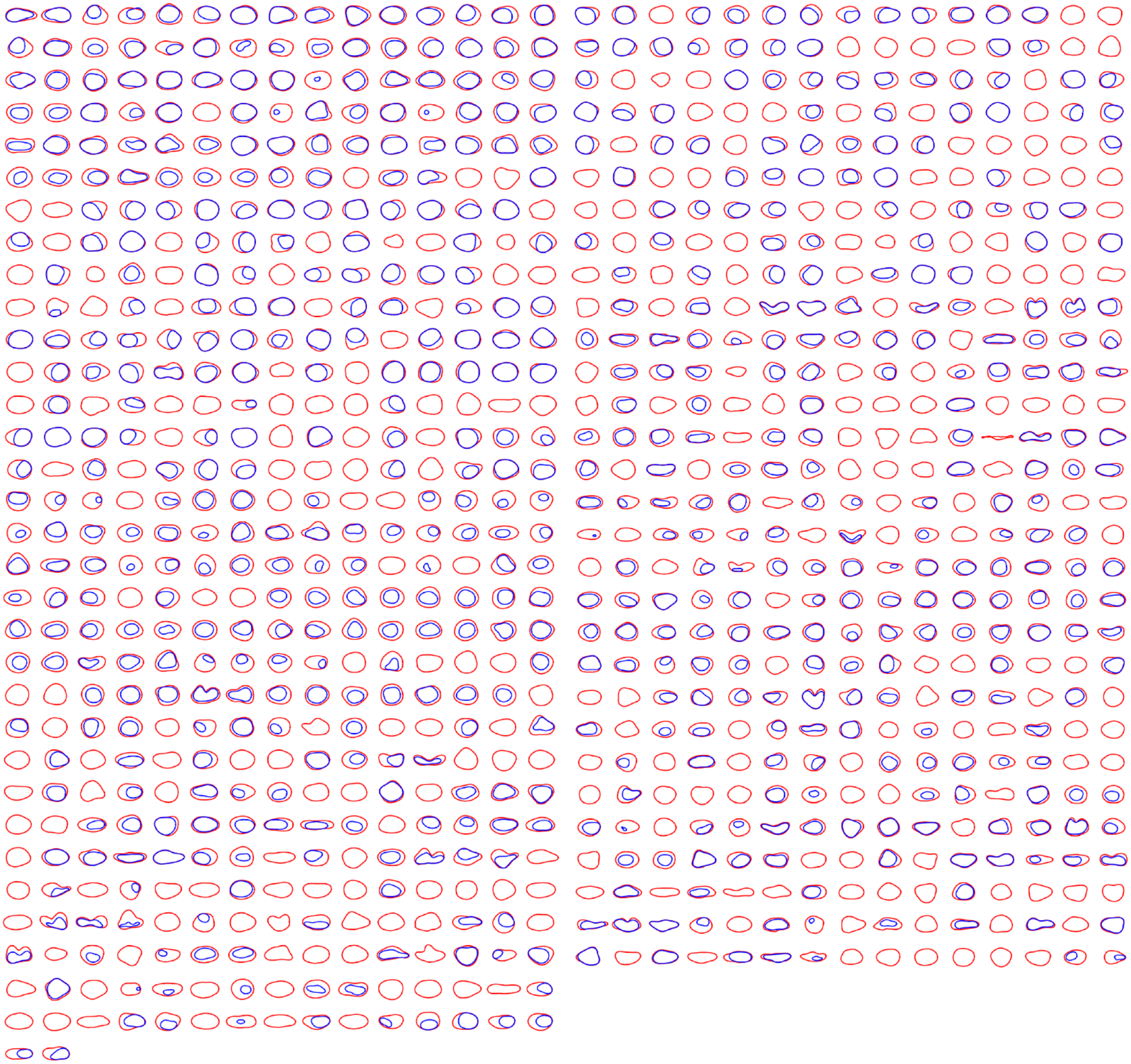

**Supplementary Fig. 3. Follicle and germinal center morphology database model using PCA decomposition.** Plot showing all follicle and germinal center shapes reconstructed from 16 PCA components analysis across the whole dataset.

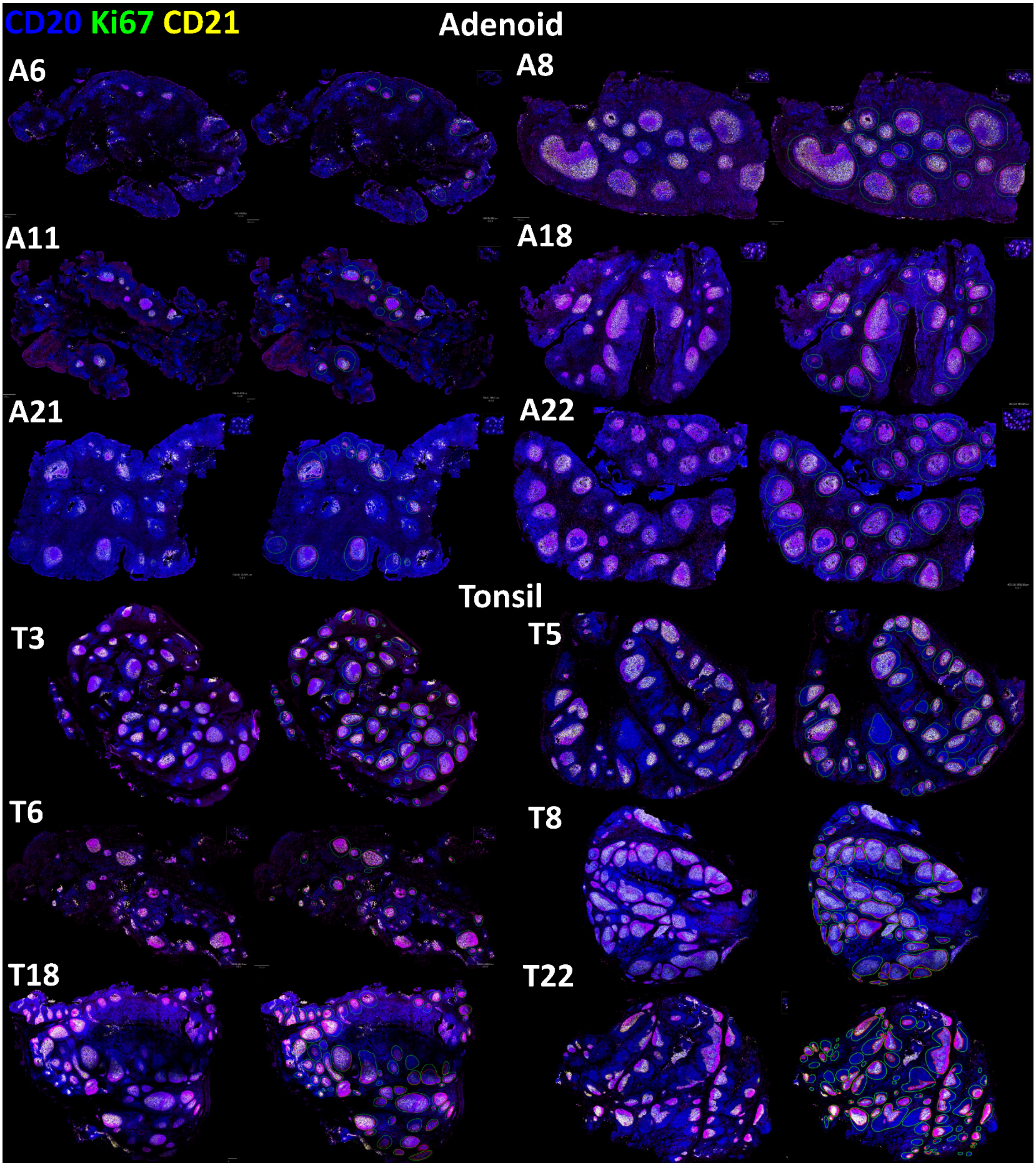
CD20 Ki67 CD21
Adenoid
A6
A8
A11
A18
A21
A22
Tonsil
T3
T5
T6
T8
T18
T22

**Supplementary Fig. 4. Multiplex imaging in human tonsil and adenoid samples.** Whole slide imaging showing CD20 (blue), Ki67 (green), and CD21 (yellow) in human adenoid (top) and tonsil (bottom) samples. Each sample is shown with only the multiplex protein markers (left) and with the segmentation result (right) of follicle regions (green contours) with their corresponding segmented germinal center regions (red contours).

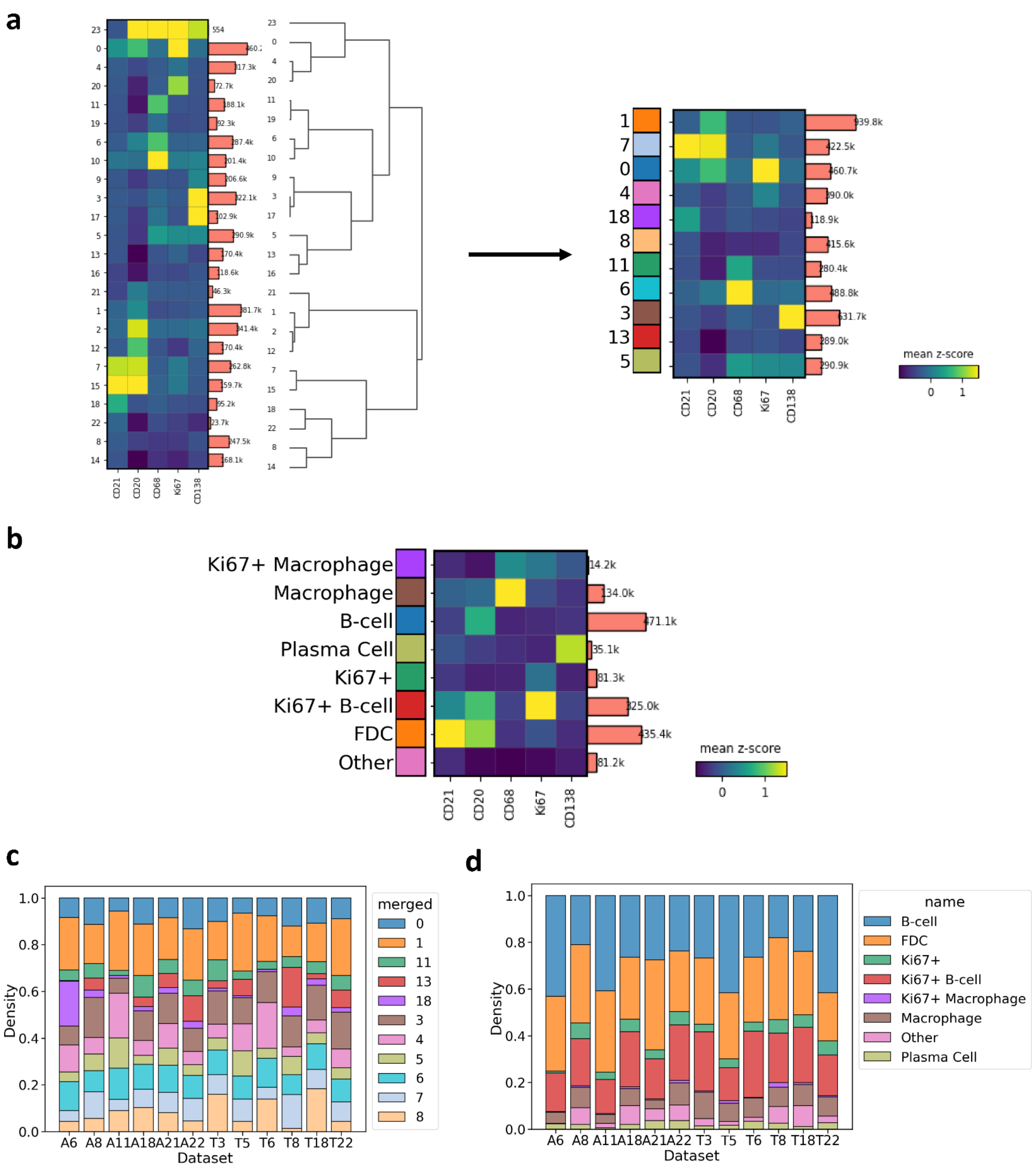

**Supplementary Fig. 5. Single cell clustering in NIH-COVID dataset.**

**a.** (Left) Heatmap showing the unsupervised clustering result using leiden clustering algorithm resulting in 24 clusters from all cells. Bar graph in the middle shows the number of cells in each cluster. Dendrogram shows the cluster similarities based on the mean expression level of each cluster. (Right) Heatmap showing the merged clustering by combining similar cluster from the left side.

**b.** Heatmap showing the unsupervised clustering annotation based on cluster mean expression in follicle regions only.

**c.** Stacked bar plot showing the distribution of clusters in all tissue regions across patient tissues.

**d.** Stacked bar plot showing the distribution of clusters in follicle regions across patient tissues.

A6

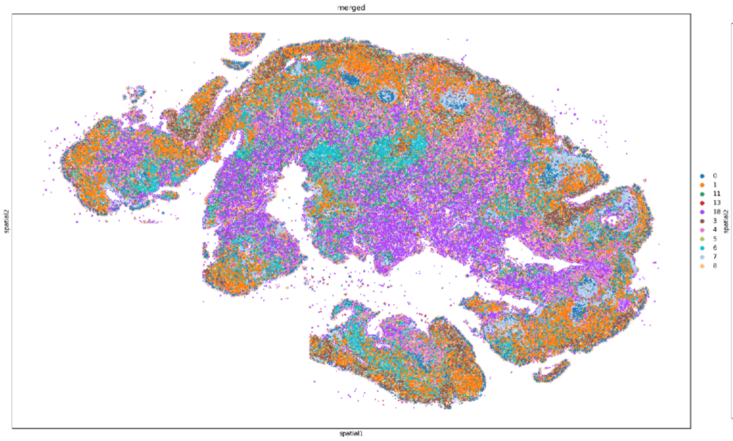

A8

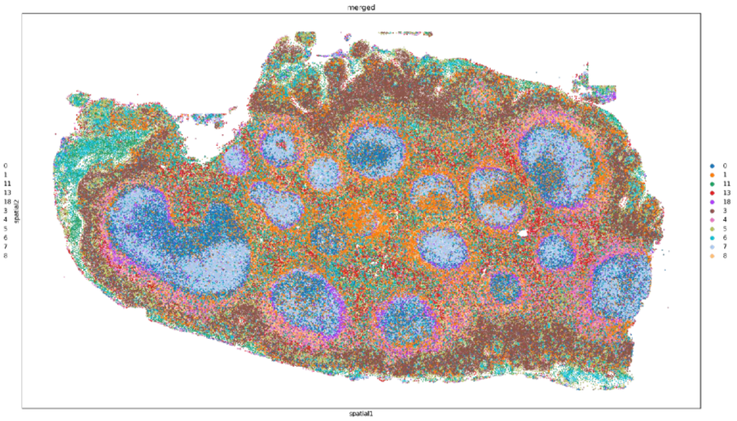

A11

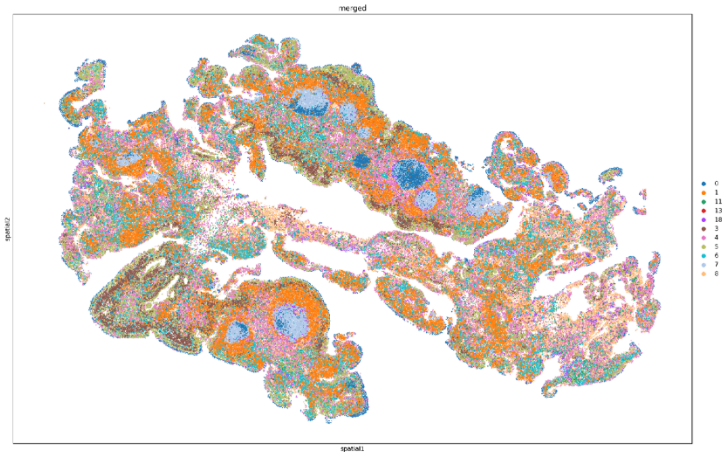

A18

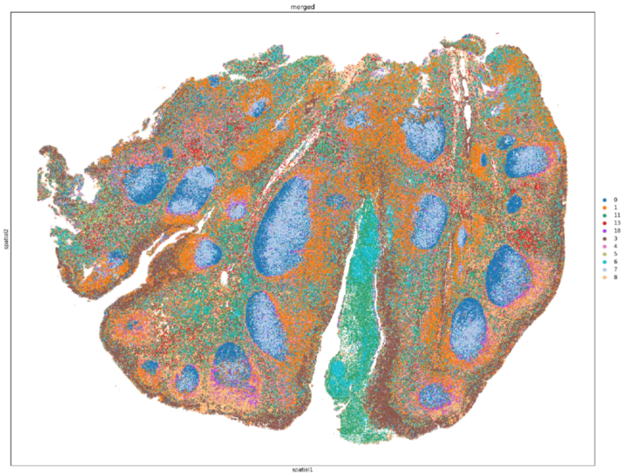

A21

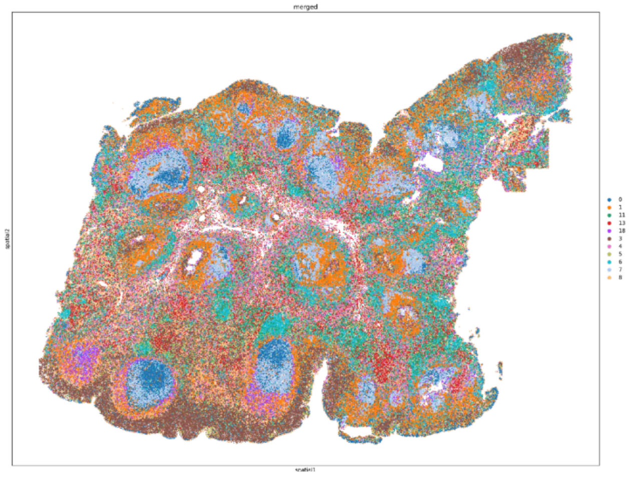

A22

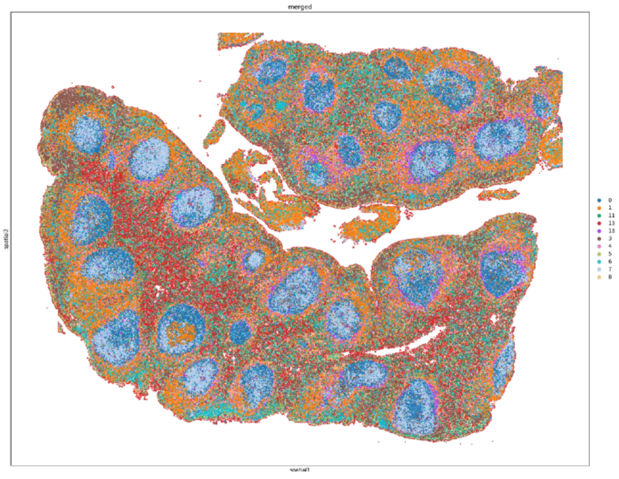

**Supplementary Fig. 6. Spatial distribution of single cell clustering in NIH-COVID dataset.** Scatter plot showing the spatial distribution of single cells in adenoid tissue samples in the NIH-COVID dataset with color corresponding to the colormap in Supplementary Fig 5. a.

T3

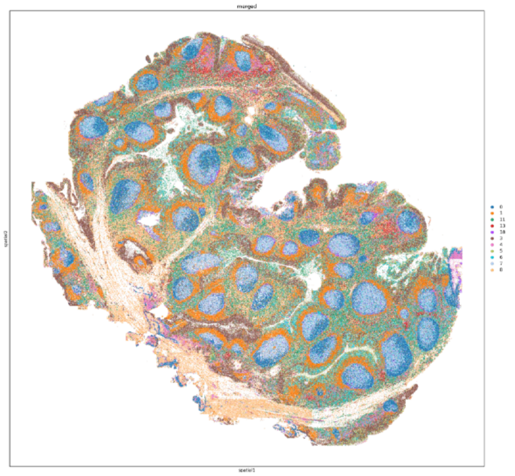

T5

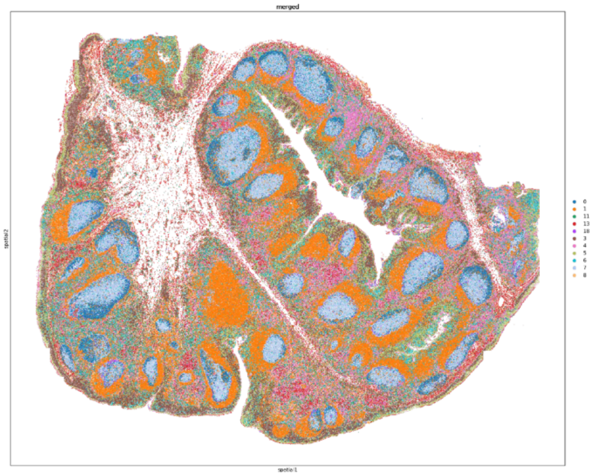

T6

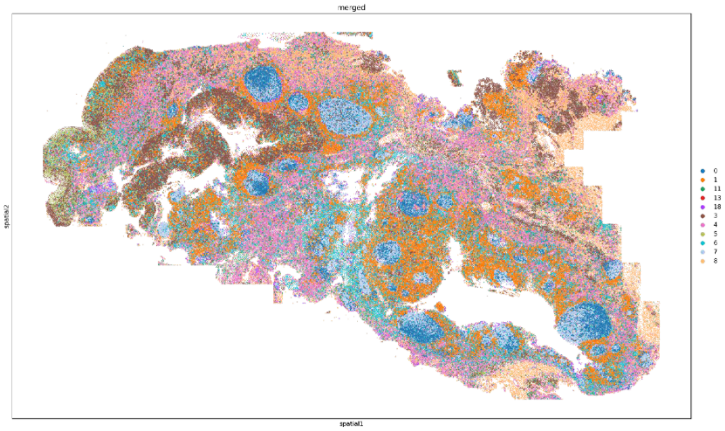

T6

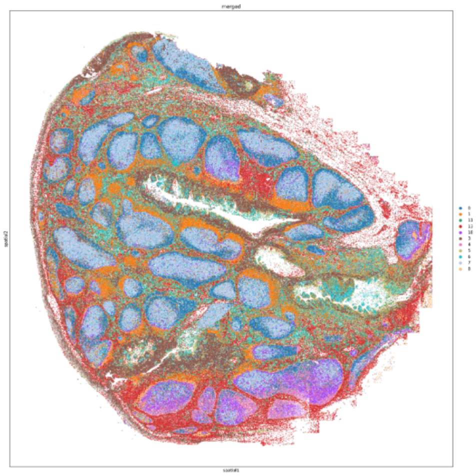

T18

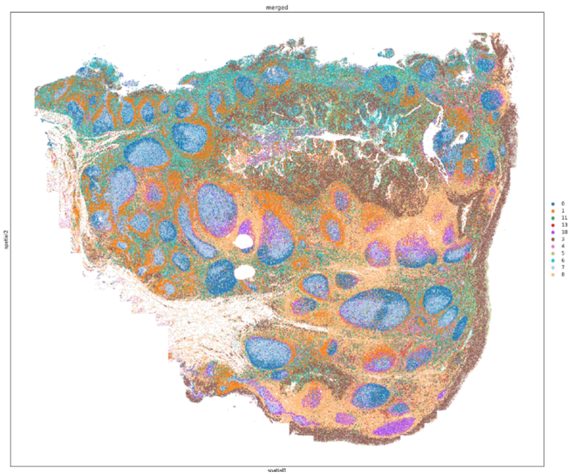

T22

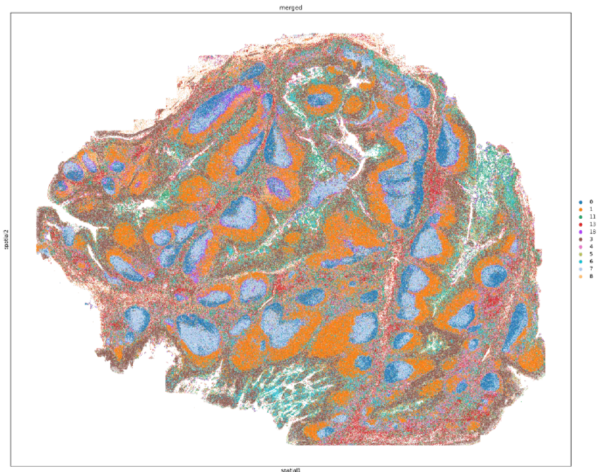

**Supplementary Fig. 7. Spatial distribution of single cell clustering in NIH-COVID dataset.** Scatter plot showing the spatial distribution of single cells tonsil tissue samples in the NIH-COVID dataset with color corresponding to the colormap in Supplementary Fig 5. a.

A6

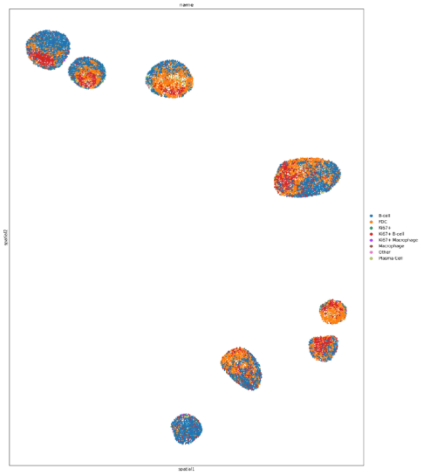

A8

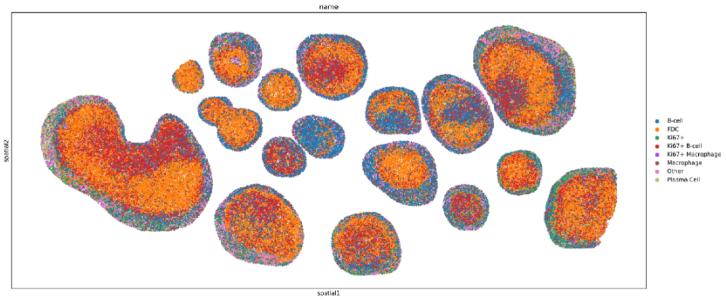

A11

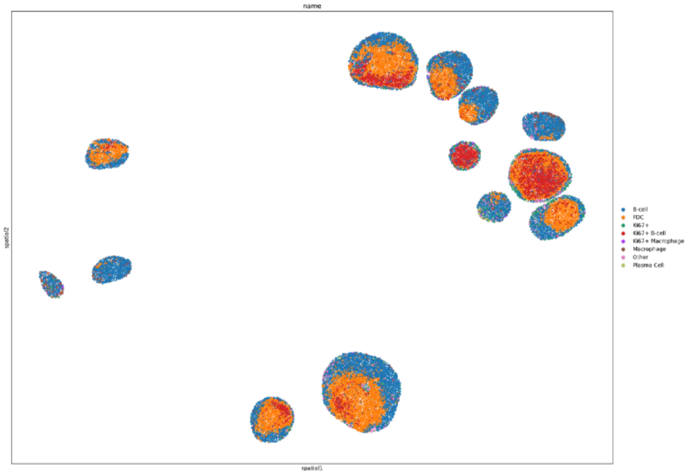

A18

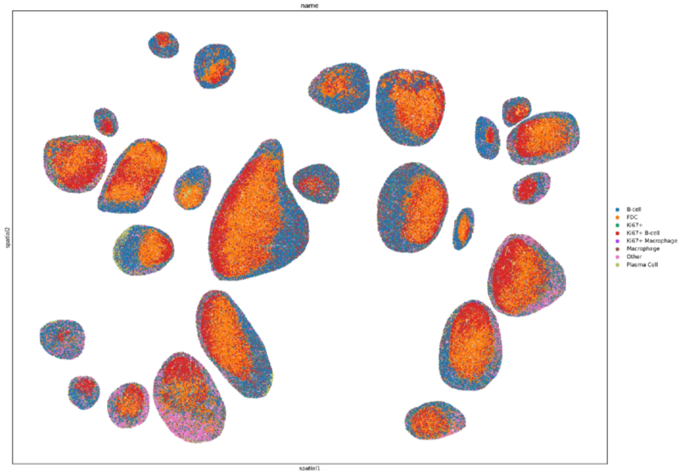

A21

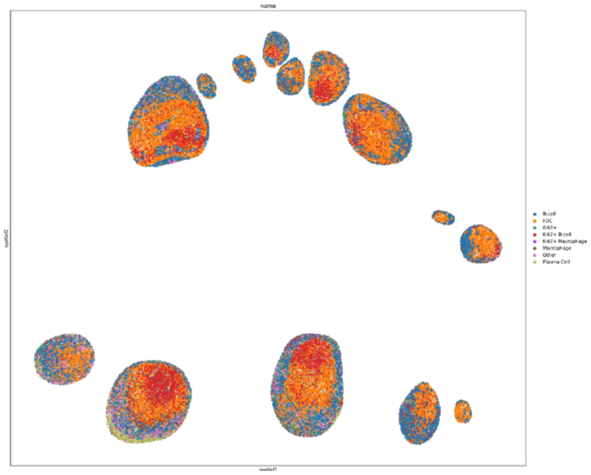

A22

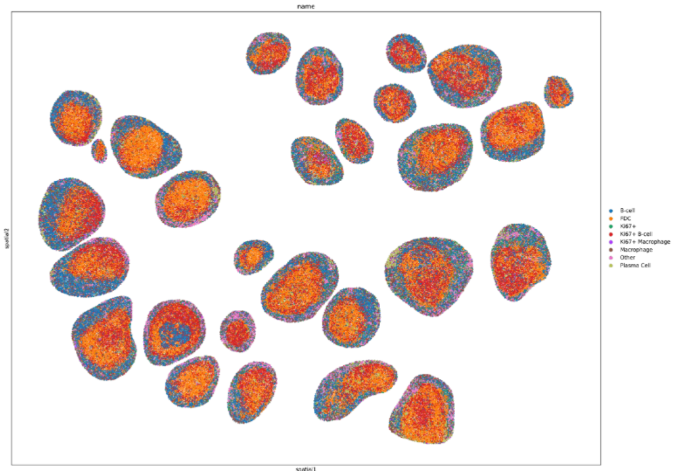

**Supplementary Fig. 8. Spatial distribution of single cell clustering in NIH-COVID dataset.** Scatter plot showing the spatial distribution of single cells inside follicle regions of adenoid tissue samples in the NIH-COVID dataset with color corresponding to the colormap in Supplementary Fig 5. b.

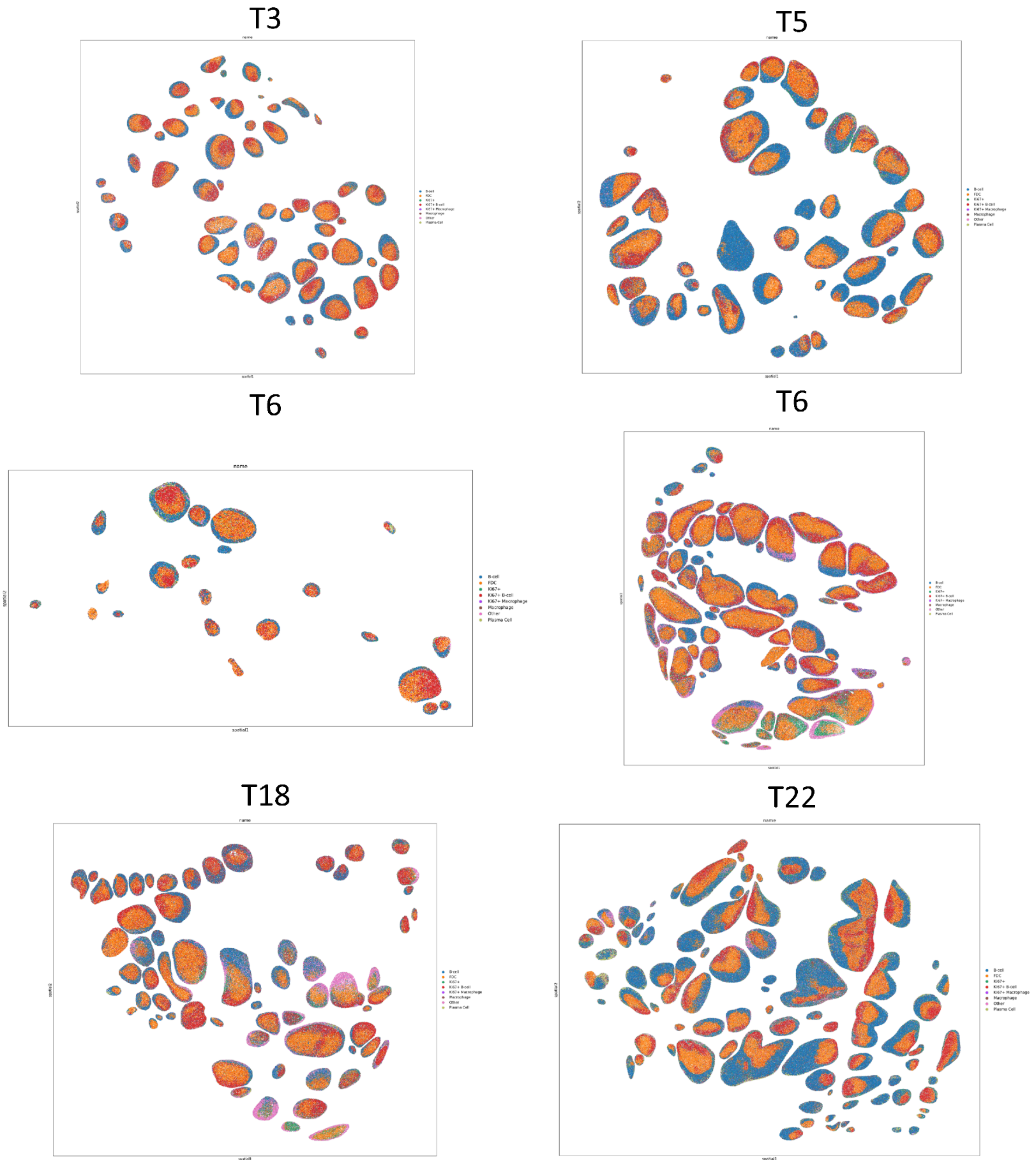
T3
T5
T6
T6
T18
T22
name
spatial1
spatial2
B-cell
FDC
Ki67+
Ki67+ B-cell
Ki67+ Macrophage
Macrophage
Other
Plasma Cell

**Supplementary Fig. 9. Spatial distribution of single cell clustering in NIH-COVID dataset.** Scatter plot showing the spatial distribution of single cells inside follicle regions of tonsil tissue samples in the NIH-COVID dataset with color corresponding to the colormap in Supplementary Fig 5. b.

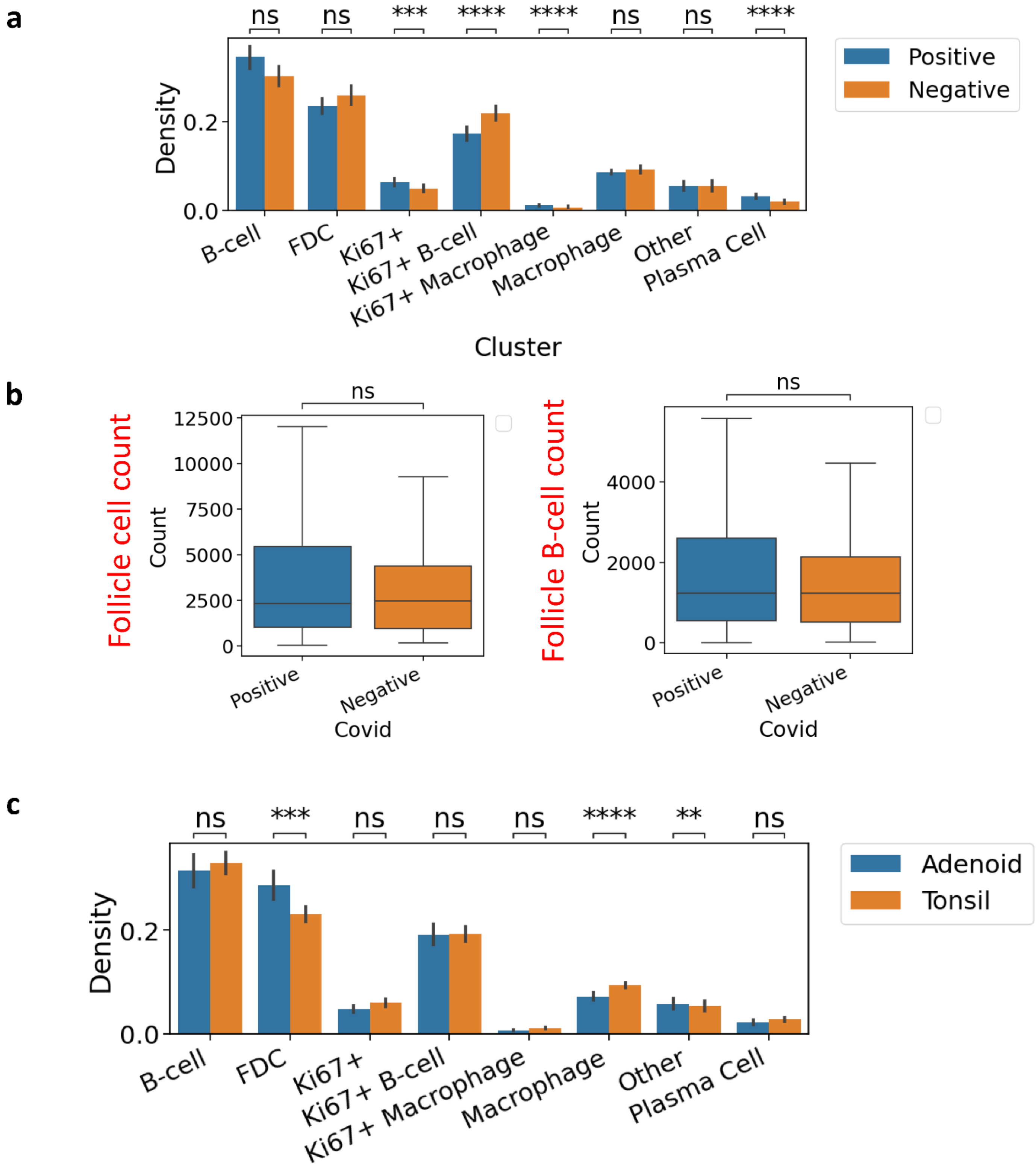
a
ns
ns
***
****
****
ns
ns
****
Density
0.2
0.0
Positive
Negative
B-cell
FDC
Ki67+
Ki67+ B-cell
Ki67+ Macrophage
Macrophage
Other
Plasma Cell
Cluster
b
ns
Follicle cell count
Count
12500
10000
7500
5000
2500
0
Positive
Negative
Covid
ns
Follicle B-cell count
Count
4000
2000
0
Positive
Negative
Covid
c
ns
***
ns
ns
ns
****
**
ns
Density
0.2
0.0
Adenoid
Tonsil
B-cell
FDC
Ki67+
Ki67+ B-cell
Ki67+ Macrophage
Macrophage
Other
Plasma Cell
Cluster

**Supplementary Fig. 10. Comparison of single-cell phenotype distribution.**

**a.** Bar plots showing the distribution comparison of single-cell phenotypes inside COVID-positive and negative follicles. Mann-Whitney-Wilcoxon test was two-sided with Bonferroni correction (ns: 0.05 < p, **: 0.001 < p <= 0.01, ***: 0.0001 < p <= 0.001, ****: p<=0.0001).

**b.** Box plots showing the distribution comparison of single-cell count inside follicles (left) and B-cells inside follicles (right) in COVID infected and control samples. Mann-Whitney-Wilcoxon test was two-sided (ns: 0.05 < p).

**c.** Bar plots showing the distribution comparison of single-cell phenotypes inside tonsil and adenoid tissue follicles. Mann-Whitney-Wilcoxon test was two-sided with Bonferroni correction (ns: 0.05 < p, **: 0.001 < p <= 0.01, ***: 0.0001 < p <= 0.001, ****: p<=0.0001).

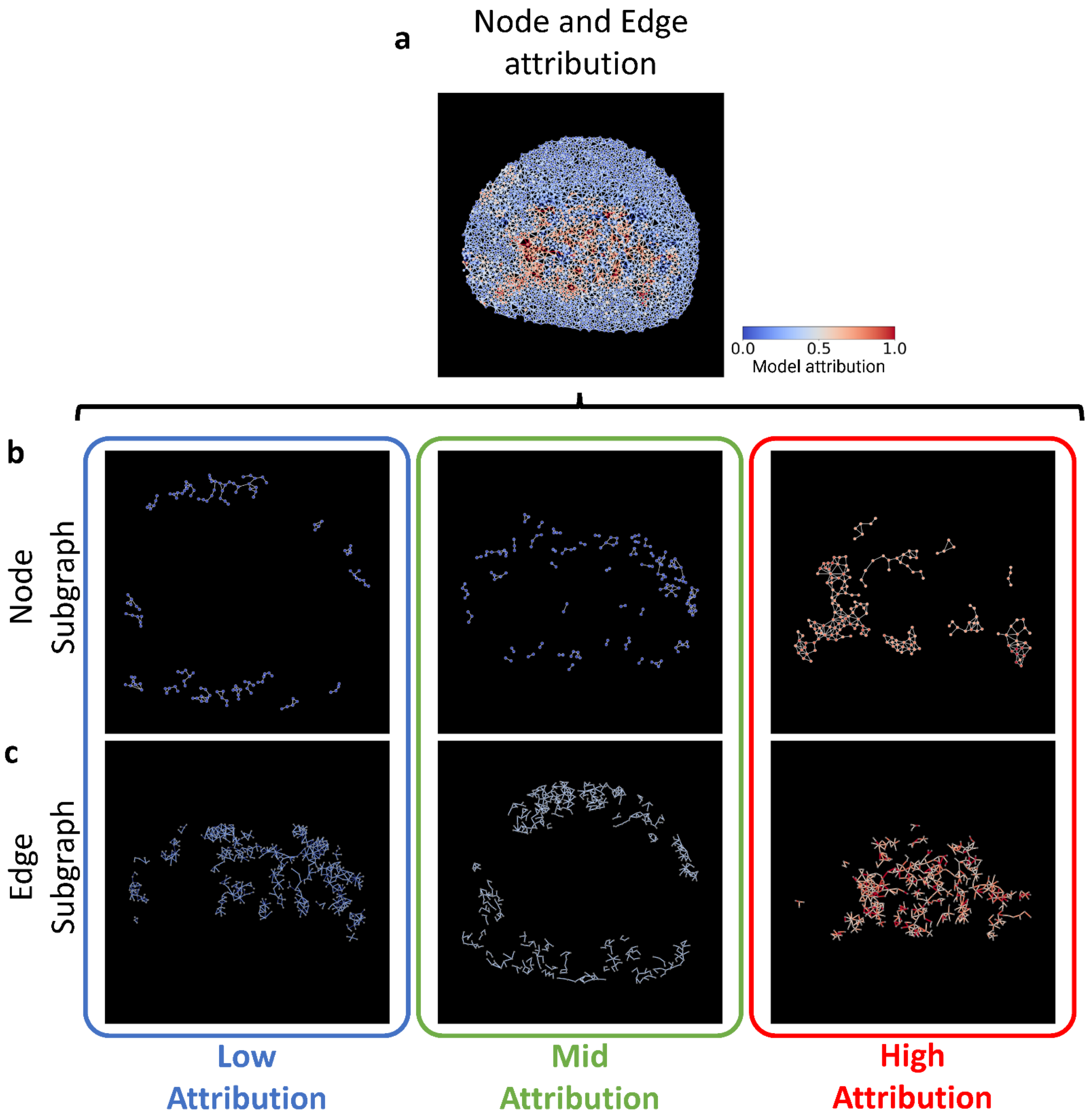
a
Node and Edge attribution
0.0
0.5
1.0
Model attribution
b
Node Subgraph
c
Edge Subgraph
Low Attribution
Mid Attribution
High Attribution

**Supplementary Fig. 11. Visual example of a COVID-positive follicle with resulting node and edge attribution and subgraph extraction.**

- **a.** A visual example of a single-cell spatial graph showing the node and edge attribution values. Each node and edge gives a color corresponding to the relative attribution value normalized at the graph level.
- **b.** A visual example of extracted low (left), medium (center), and high (right) attribution subgraphs using node-level attribution scores. From the node level attribution scores, the lower 10 percentile, middle 10 percentile, and top 10 percentile nodes are extracted and the edges connecting these nodes are maintained while the rest of the edges are deleted. Individual nodes without connecting nodes are also deleted. This provides the SNWOFLAKE pipeline node attribution-induced subgraphs.
- **c.** A visual example of extracted low (left), medium (center), and high (right) attribution subgraphs using edge-level attribution scores. From the edge level attribution scores, the lower 10 percentile, middle 10 percentile, and top 10 percentile edges are extracted and the nodes at the extremities are maintained while the rest of the nodes are deleted. Individual edges without connecting components are also deleted. This provides the SNWOFLAKE pipeline edge attribution-induced subgraphs.

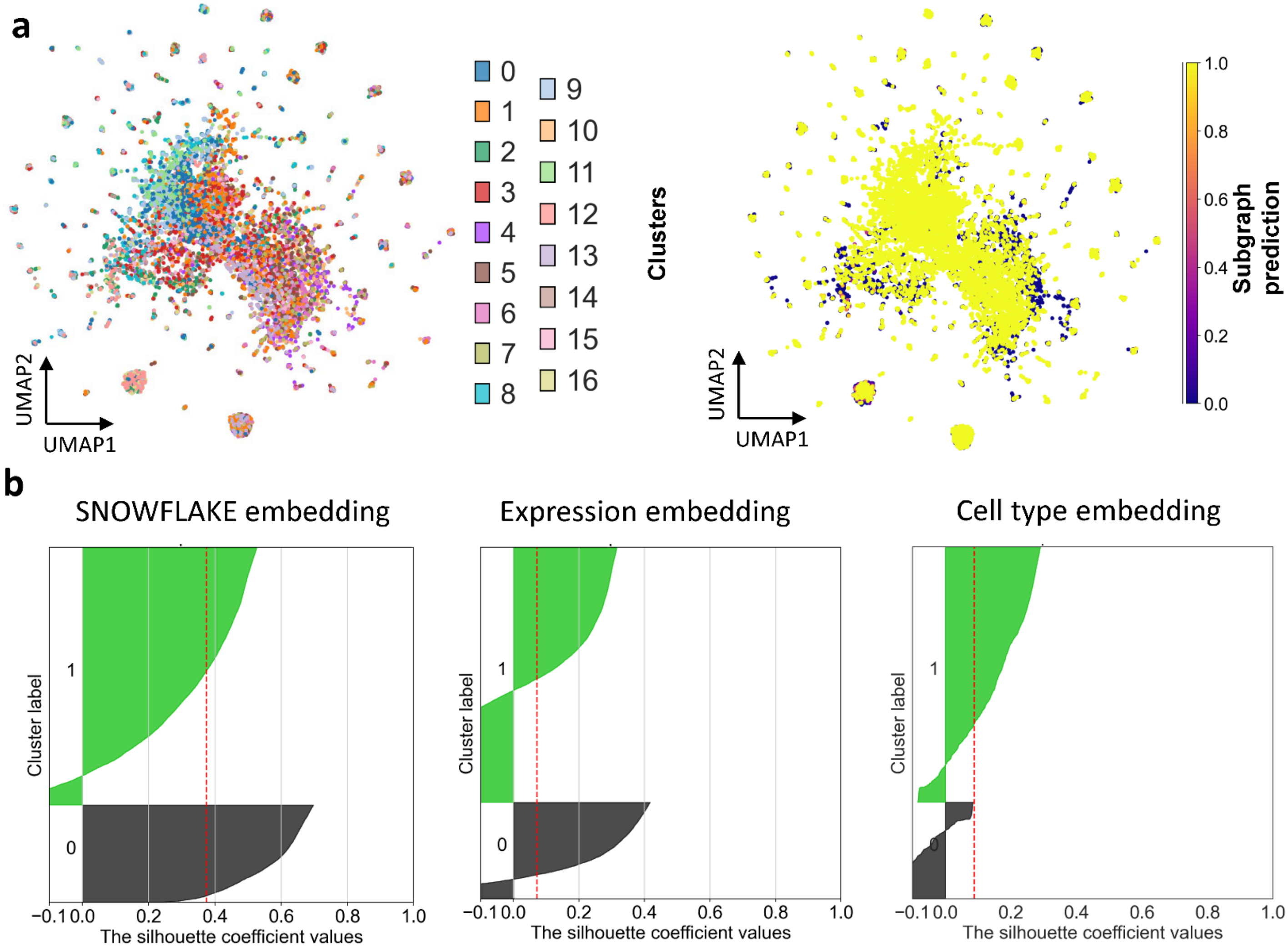
a
UMAP2
UMAP1
Clusters
Subgraph prediction
1.0
0.8
0.6
0.4
0.2
0.0
b
SNOWFLAKE embedding
Expression embedding
Cell type embedding
Cluster label
The silhouette coefficient values
−0.1
0.0
0.2
0.4
0.6
0.8
1.0

**Supplementary Fig. 12. Comparison of subgraph clustering using cell expression level and cell type density.**

**a.** UMAPs show the SC embedding low dimensional projection using single-cell type densities instead of subgraph embedding obtained from the SNOWFLAKE pipeline. The UMAPs show no separation between SC clusters nor positive and negative predictive SC.

**b.** Silhouette plot showing how close each point in one cluster is to points in the neighboring clusters. The silhouette plots show the relative size of the COVID positive and negative follicles and the corresponding silhouette score of each case. A higher silhouette score means better separation between conditions. Three silhouette plots showing the silhouette score for UMAP embedding using subgraph SNOWFLAKE embeddings (left), subgraph mean marker expressions (left), and subgraph single-cell phenotype densities (right).

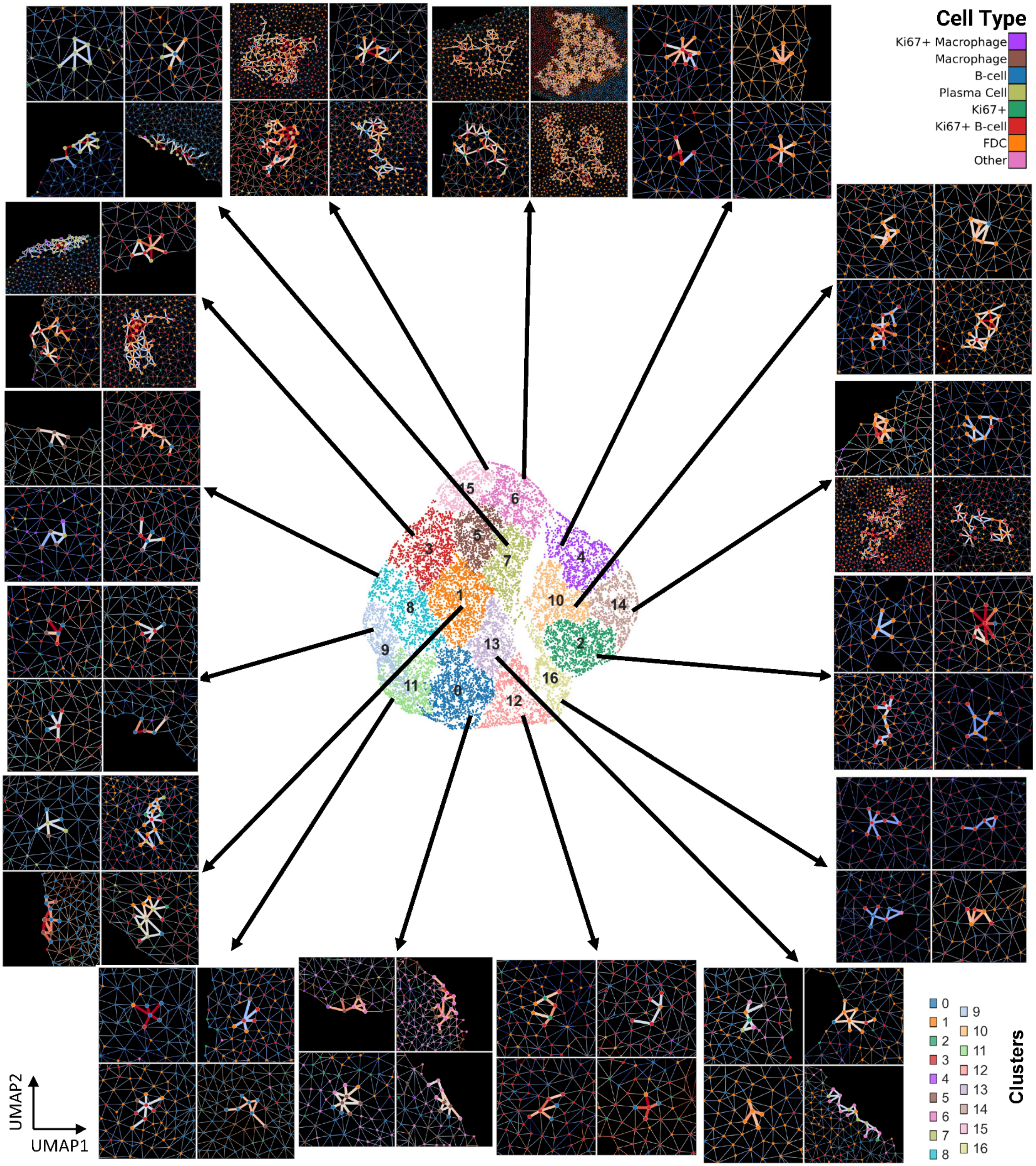
Cell Type
Ki67+ Macrophage
Macrophage
B-cell
Plasma Cell
Ki67+
Ki67+ B-cell
FDC
Other
15
6
3
5
7
4
1
8
10
14
9
13
2
11
0
16
12
UMAP2
UMAP1
Clusters

**Supplementary Fig. 13. Visual representations of the latent space distribution of the subgraph clusters obtained from SNOWFLAKE embedding.** The scatter plot in the center shows the low dimensional UMAP embedding of the subgraph with the color corresponding to distinct subgraph clusters. Each arrow shows four visual examples of subgraphs close to each subgraph cluster centroid region. The node color represents distinct cell types and the edge color represents the relative attribution score for each edge from the SNOWFLAKE pipeline.

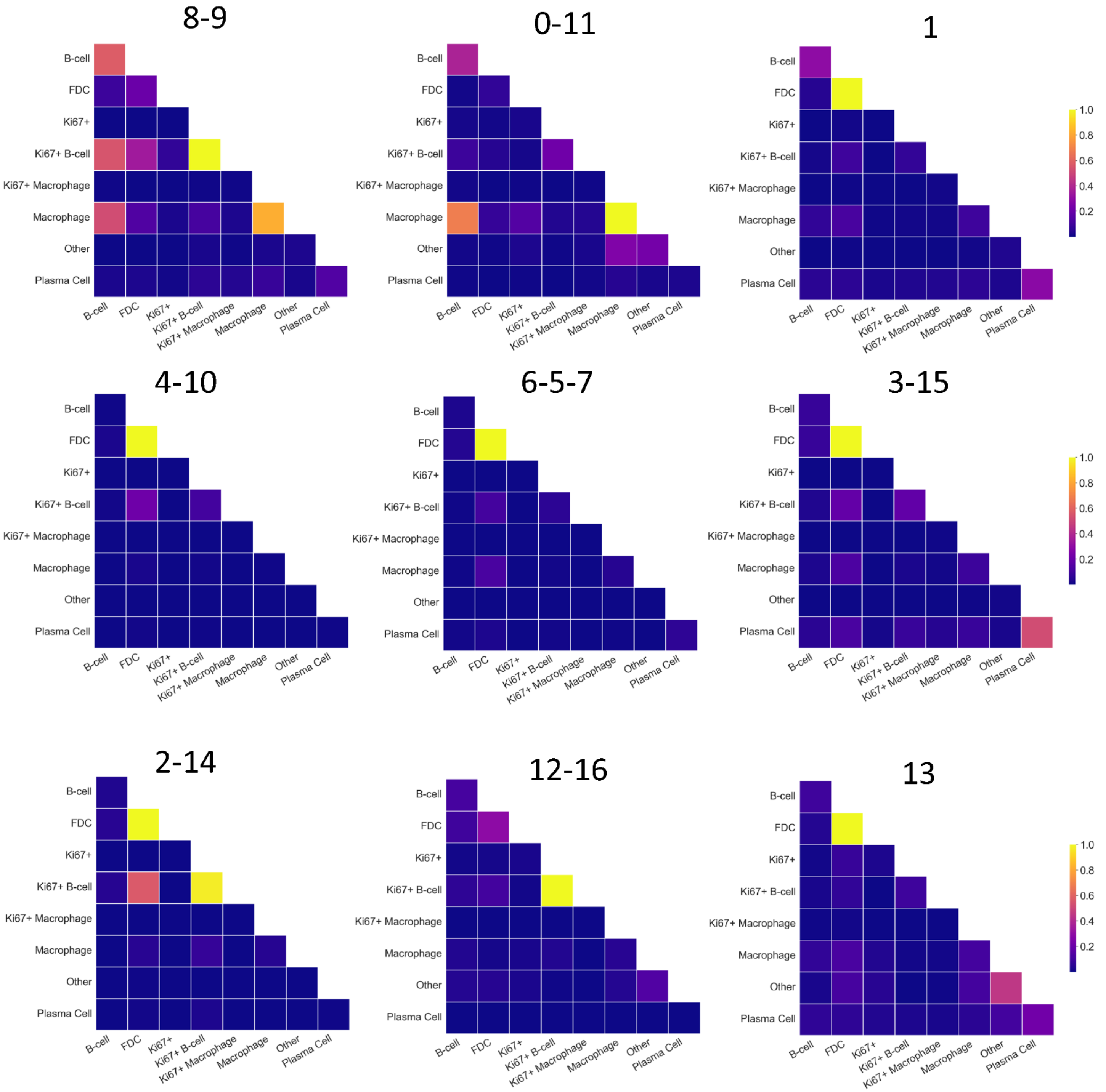
8-9
0-11
1
4-10
6-5-7
3-15
2-14
12-16
13
B-cell
FDC
Ki67+
Ki67+ B-cell
Ki67+ Macrophage
Macrophage
Other
Plasma Cell
1.0
0.8
0.6
0.4
0.2

**Supplementary Fig. 14. Edge connectivity heatmaps across all subgraph clusters.** Heatmap showing the cell-type level connectivity densities for subgraph clusters obtained from SNOWFLAKE embeddings. Subgraph clusters are grouped based on their similarity from dendrogram clustering.

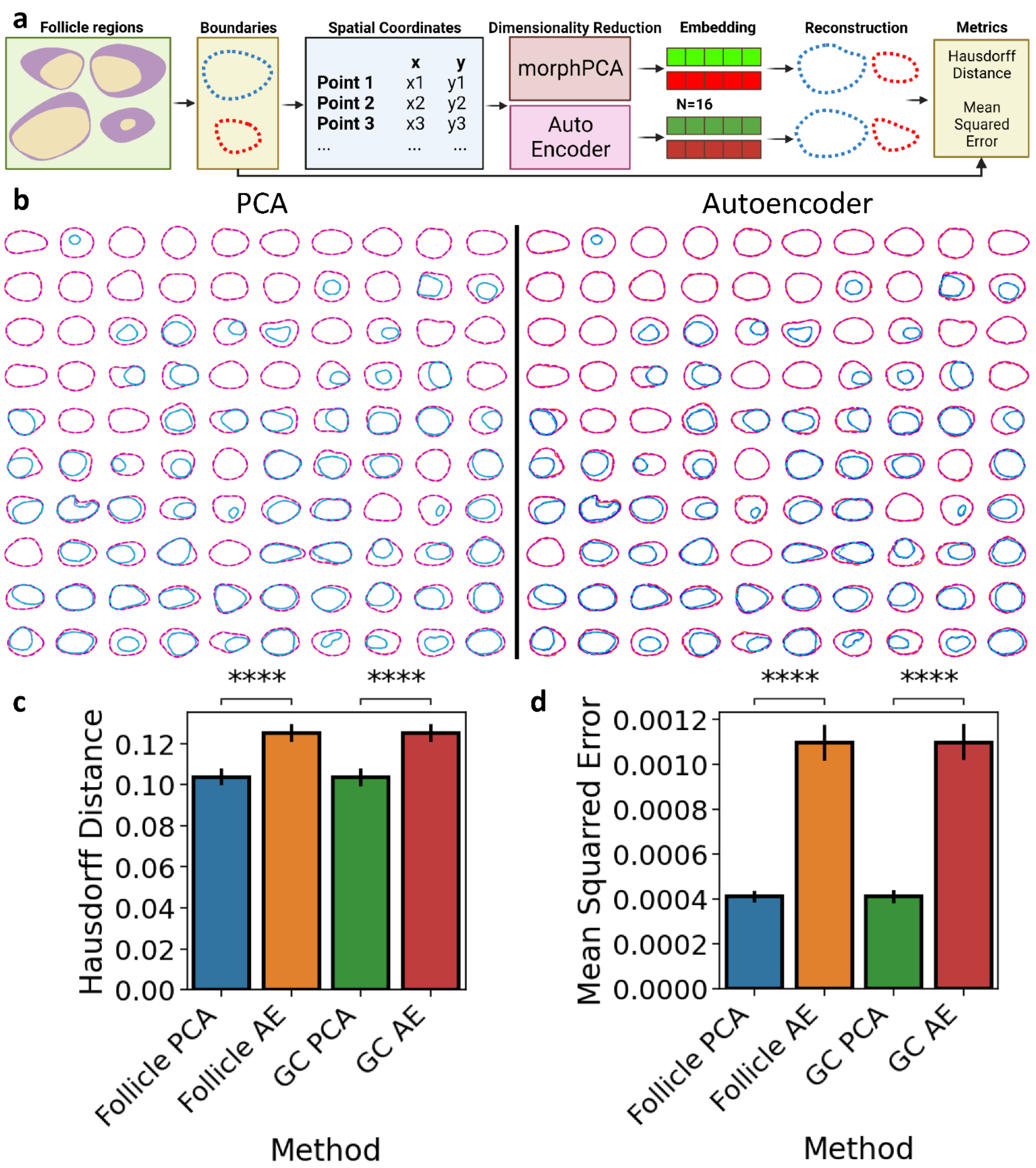

a
Follicle regions
Boundaries
Spatial Coordinates
x y
Point 1 x1 y1
Point 2 x2 y2
Point 3 x3 y3
Dimensionality Reduction
morphPCA
Auto Encoder
Embedding
N=16
Reconstruction
Metrics
Hausdorff Distance
Mean Squared Error
b
PCA
Autoencoder
c
Hausdorff Distance
Method
Follicle PCA
Follicle AE
GC PCA
GC AE
****
d
Mean Squarred Error
Method
Follicle PCA
Follicle AE
GC PCA
GC AE
****

**Supplementary Fig. 15. Comparison of morphology analysis between PCA and Autoencoder methods.**

- **a.** Schematic showing the comparison of morphology feature extraction between PCA and Autoencoder models. From normalized and aligned follicle and germinal center boundaries, the spatial coordinates are sampled at equal space distances. Low dimensional embeddings (n=16) are obtained for follicle and germinal center coordinates using PCA and Autoencoder model. The reconstruction of boundary coordinates are compared with the original boundary coordinates using Hausdorff distance and Mean Squared Error.
- **b.** Example of reconstructed (in dashed lines) follicles and germinal center boundaries compared to original boundaries (in full lines) for the PCA methods (left) and Autoencoder methods (right).
- **c.** Bar plot showing the average Hausdorff distance (left) and Mean Squared Error (right) comparison of follicle and germinal center boundaries reconstruction using low dimensional embeddings (n=16) from PCA and Autoencoder models. Mann-Whitney-Wilcoxon test was two-sided with Bonferroni correction (ns: $0.05 < p$, ****: $p<=0.0001$).

### Supplementary Tables

| N | Reference | Modality | Tissue ID | Tissue Origin | GC count | Follicle count | Cell count |
|---|---|---|---|---|---|---|---|
| 1 | (*1*) | CODEX | Tonsil_1 | Tonsil | 85 | 88 | 543918 |
| 2 | (*2*) | CycIF | Tonsil_1 | Tonsil | 188 | 269 | 1272090 |
| 3 | (*3*) | CycIF | Tonsil_1 | Tonsil | 89 | 91 | 2067691 |
| 4 | (*4*) | CODEX | Tonsil_CNMC3 | Tonsil | 52 | 56 | 586504 |
| | | | Tonsil_CNMC5 | Tonsil | 41 | 45 | 495603 |
| | | | Tonsil_CNMC6 | Tonsil | 19 | 22 | 245888 |
| | | | Tonsil_CNMC8 | Tonsil | 64 | 67 | 501731 |
| | | | Tonsil_CNMC18 | Tonsil | 61 | 64 | 644920 |
| | | | Tonsil_CNMC22 | Tonsil | 60 | 83 | 873477 |
| | | | Adenoid_CNMC6 | Adenoid | 6 | 8 | 124376 |
| | | | Adenoid_CNMC8 | Adenoid | 17 | 17 | 187420 |
| | | | Adenoid_CNMC11 | Adenoid | 9 | 13 | 160322 |
| | | | Adenoid_CNMC18 | Adenoid | 25 | 25 | 411600 |
| | | | Adenoid_CNMC21 | Adenoid | 9 | 14 | 227886 |
| | | | Adenoid_CNMC22 | Adenoid | 28 | 28 | 269532 |
| 5 | (*5*) | CycIF | Tonsil | Tonsil | 22 | 40 | 266791 |
| Total | | | | | 775 | 930 | 8879749 |

**Supplementary Table 1. Statistical summary of the datasets curated in this study for morphology analysis.**

| Model | Type | Accuracy | AUC | F1 |
|---|---|---|---|---|
| Adaboost | Mean Expression | 0.765 ± 0.03 | 0.768 ± 0.056 | 0.416 ± 0.108 |
| | Morph PCA | 0.715 ± 0.017 | 0.677 ± 0.033 | 0.333 ± 0.107 |
| | Mean Expression + Morph PCA | 0.767 ± 0.043 | **0.808 ± 0.031** | 0.476 ± 0.062 |
| DecisionTree | Mean Expression | 0.751 ± 0.007 | 0.661 ± 0.035 | 0.477 ± 0.056 |
| | Morph PCA | 0.661 ± 0.07 | 0.531 ± 0.065 | 0.29 ± 0.08 |
| | Mean Expression + Morph PCA | 0.74 ± 0.049 | **0.675 ± 0.059** | 0.497 ± 0.07 |
| GradientBoosting | Mean Expression | 0.817 ± 0.026 | 0.828 ± 0.03 | 0.513 ± 0.103 |
| | Morph PCA | 0.742 ± 0.041 | 0.67 ± 0.042 | 0.25 ± 0.086 |
| | Mean Expression + Morph PCA | 0.817 ± 0.052 | **0.844 ± 0.051** | 0.554 ± 0.08 |
| NaiveBayes | Mean Expression | 0.774 ± 0.025 | 0.73 ± 0.035 | 0.268 ± 0.094 |
| | Morph PCA | 0.613 ± 0.035 | 0.684 ± 0.074 | 0.468 ± 0.061 |
| | Mean Expression + Morph PCA | 0.663 ± 0.051 | **0.74 ± 0.084** | 0.504 ± 0.07 |
| RandomForest | Mean Expression | 0.812 ± 0.032 | **0.85 ± 0.038** | 0.502 ± 0.052 |
| | Morph PCA | 0.758 ± 0.03 | 0.619 ± 0.056 | 0.034 ± 0.048 |
| | Mean Expression + Morph PCA | 0.79 ± 0.033 | 0.798 ± 0.052 | 0.246 ± 0.065 |
| SVM | Mean Expression | 0.806 ± 0.028 | **0.835 ± 0.035** | 0.387 ± 0.067 |
| | Morph PCA | 0.763 ± 0.03 | 0.624 ± 0.048 | 0.0 ± 0.0 |
| | Mean Expression + Morph PCA | 0.769 ± 0.032 | 0.774 ± 0.03 | 0.056 ± 0.053 |
| MLP | Mean Expression | 0.79 ± 0.036 | **0.748 ± 0.045** | 0.268 ± 0.108 |
| | Morph PCA | 0.753 ± 0.034 | 0.657 ± 0.085 | 0.08 ± 0.075 |
| | Mean Expression + Morph PCA | 0.757 ± 0.032 | 0.741 ± 0.059 | 0.04 ± 0.047 |

**Supplementary Table 2. Machine learning and multi-instance learning models benchmarked based on the mean expression, morphology, and combination of morphology and mean expression from follicles.**

| Model | Pooling Strategy | Morphology Fusion | Accuracy | AUC | F1 |
|---|---|---|---|---|---|
| MLP | Gated Attention | Concatenation | 0.821 ± 0.046 | 0.9 ± 0.035 | 0.85 ± 0.04 |
| | | Expression Only | 0.609 ± 0.101 | 0.879 ± 0.033 | 0.486 ± 0.233 |
| | | Tensor Decomposition | 0.815 ± 0.029 | **0.915 ± 0.027** | 0.857 ± 0.02 |
| | Global Attention | Concatenation | 0.764 ± 0.034 | 0.903 ± 0.045 | 0.76 ± 0.05 |
| | | Expression Only | 0.756 ± 0.055 | **0.896 ± 0.042** | 0.747 ± 0.068 |
| | | Tensor Decomposition | 0.734 ± 0.081 | 0.891 ± 0.032 | 0.709 ± 0.134 |
| | Max | Concatenation | 0.722 ± 0.057 | 0.852 ± 0.061 | 0.752 ± 0.09 |
| | | Expression Only | 0.636 ± 0.106 | 0.856 ± 0.075 | 0.558 ± 0.192 |
| | | Tensor Decomposition | 0.65 ± 0.17 | **0.894 ± 0.046** | 0.516 ± 0.368 |
| | Mean | Concatenation | 0.785 ± 0.042 | **0.909 ± 0.035** | 0.782 ± 0.06 |
| | | Expression Only | 0.803 ± 0.027 | 0.904 ± 0.03 | 0.813 ± 0.029 |
| | | Tensor Decomposition | 0.824 ± 0.038 | 0.896 ± 0.034 | 0.847 ± 0.041 |
| GAT | Gated Attention | Concatenation | 0.833 ± 0.102 | 0.937 ± 0.062 | 0.875 ± 0.064 |
| | | Expression Only | 0.884 ± 0.016 | 0.953 ± 0.013 | 0.9 ± 0.013 |
| | | Tensor Decomposition | 0.842 ± 0.078 | **0.957 ± 0.031** | 0.879 ± 0.054 |
| | Global Attention | Concatenation | 0.821 ± 0.081 | 0.937 ± 0.029 | 0.821 ± 0.111 |
| | | Expression Only | 0.777 ± 0.05 | 0.932 ± 0.037 | 0.771 ± 0.068 |
| | | Tensor Decomposition | 0.85 ± 0.061 | **0.937 ± 0.038** | 0.882 ± 0.042 |
| | Max | Concatenation | 0.728 ± 0.084 | 0.885 ± 0.043 | 0.806 ± 0.043 |
| | | Expression Only | 0.83 ± 0.103 | **0.96 ± 0.031** | 0.872 ± 0.062 |
| | | Tensor Decomposition | 0.841 ± 0.04 | 0.878 ± 0.053 | 0.865 ± 0.037 |
| | Mean | Concatenation | 0.854 ± 0.075 | 0.95 ± 0.036 | 0.884 ± 0.054 |
| | | Expression Only | 0.863 ± 0.082 | 0.958 ± 0.037 | 0.892 ± 0.059 |
| | | Tensor Decomposition | 0.926 ± 0.01 | **0.973 ± 0.009** | 0.938 ± 0.008 |
| GCN | Gated Attention | Concatenation | 0.759 ± 0.048 | 0.837 ± 0.072 | 0.78 ± 0.044 |
| | | Expression Only | 0.794 ± 0.037 | 0.86 ± 0.068 | 0.821 ± 0.031 |
| | | Tensor Decomposition | 0.779 ± 0.029 | **0.861 ± 0.031** | 0.805 ± 0.032 |
| | Global Attention | Concatenation | 0.746 ± 0.042 | **0.833 ± 0.062** | 0.758 ± 0.047 |
| | | Expression Only | 0.744 ± 0.036 | 0.792 ± 0.065 | 0.78 ± 0.034 |
| | | Tensor Decomposition | 0.755 ± 0.06 | 0.819 ± 0.068 | 0.781 ± 0.059 |
| | Max | Concatenation | 0.797 ± 0.061 | 0.86 ± 0.035 | 0.82 ± 0.067 |
| | | Expression Only | 0.77 ± 0.041 | 0.885 ± 0.033 | 0.816 ± 0.031 |
| | | Tensor Decomposition | 0.827 ± 0.025 | **0.913 ± 0.052** | 0.856 ± 0.024 |
| | Mean | Concatenation | 0.74 ± 0.062 | 0.826 ± 0.051 | 0.771 ± 0.072 |
| | | Expression Only | 0.759 ± 0.052 | 0.817 ± 0.082 | 0.797 ± 0.041 |
| | | Tensor Decomposition | 0.765 ± 0.041 | **0.834 ± 0.053** | 0.791 ± 0.044 |
| GINConv | Gated Attention | Concatenation | 0.833 ± 0.035 | 0.886 ± 0.024 | 0.858 ± 0.032 |
| | | Expression Only | 0.803 ± 0.046 | 0.885 ± 0.016 | 0.828 ± 0.035 |
| | | Tensor Decomposition | 0.802 ± 0.04 | **0.888 ± 0.029** | 0.838 ± 0.029 |
| | Global Attention | Concatenation | 0.803 ± 0.019 | 0.865 ± 0.036 | 0.831 ± 0.019 |
| | | Expression Only | 0.798 ± 0.038 | 0.873 ± 0.033 | 0.821 ± 0.039 |
| | | Tensor Decomposition | 0.808 ± 0.042 | **0.885 ± 0.032** | 0.824 ± 0.047 |
| | Max | Concatenation | 0.785 ± 0.019 | 0.854 ± 0.035 | 0.803 ± 0.019 |
| | | Expression Only | 0.795 ± 0.025 | **0.893 ± 0.017** | 0.82 ± 0.028 |
| | | Tensor Decomposition | 0.815 ± 0.009 | 0.875 ± 0.024 | 0.848 ± 0.013 |
| | Mean | Concatenation | 0.82 ± 0.013 | 0.882 ± 0.024 | 0.846 ± 0.012 |
| | | Expression Only | 0.821 ± 0.028 | 0.881 ± 0.02 | 0.845 ± 0.026 |
| | | Tensor Decomposition | 0.841 ± 0.027 | **0.892 ± 0.024** | 0.863 ± 0.02 |
| GraphConv | | Concatenation | 0.777 ± 0.029 | 0.867 ± 0.028 | 0.788 ± 0.033 |

| | | | | | |
|---|---|---|---|---|---|
| | Gated Attention | Expression Only | 0.835 ± 0.046 | **0.897 ± 0.044** | 0.859 ± 0.043 |
| | | Tensor Decomposition | 0.776 ± 0.037 | 0.863 ± 0.038 | 0.813 ± 0.034 |
| | Global Attention | Concatenation | 0.762 ± 0.039 | 0.849 ± 0.043 | 0.783 ± 0.047 |
| | | Expression Only | 0.795 ± 0.039 | **0.853 ± 0.035** | 0.824 ± 0.031 |
| | | Tensor Decomposition | 0.771 ± 0.045 | 0.842 ± 0.029 | 0.815 ± 0.032 |
| | Max | Concatenation | 0.786 ± 0.026 | 0.848 ± 0.044 | 0.809 ± 0.026 |
| | | Expression Only | 0.842 ± 0.017 | **0.927 ± 0.026** | 0.855 ± 0.013 |
| | | Tensor Decomposition | 0.814 ± 0.044 | 0.909 ± 0.037 | 0.833 ± 0.043 |
| | Mean | Concatenation | 0.764 ± 0.035 | 0.853 ± 0.048 | 0.785 ± 0.032 |
| | | Expression Only | 0.798 ± 0.046 | 0.846 ± 0.045 | 0.829 ± 0.041 |
| | | Tensor Decomposition | 0.783 ± 0.032 | **0.867 ± 0.032** | 0.815 ± 0.04 |
| SAGEConv | Gated Attention | Concatenation | 0.842 ± 0.07 | **0.932 ± 0.025** | 0.877 ± 0.047 |
| | | Expression Only | 0.838 ± 0.034 | 0.904 ± 0.036 | 0.859 ± 0.04 |
| | | Tensor Decomposition | 0.827 ± 0.052 | 0.899 ± 0.062 | 0.856 ± 0.05 |
| | Global Attention | Concatenation | 0.808 ± 0.033 | 0.897 ± 0.045 | 0.826 ± 0.032 |
| | | Expression Only | 0.877 ± 0.027 | **0.933 ± 0.02** | 0.889 ± 0.028 |
| | | Tensor Decomposition | 0.835 ± 0.062 | 0.867 ± 0.069 | 0.864 ± 0.05 |
| | Max | Concatenation | 0.788 ± 0.079 | 0.89 ± 0.066 | 0.811 ± 0.063 |
| | | Expression Only | 0.91 ± 0.04 | **0.964 ± 0.033** | 0.924 ± 0.031 |
| | | Tensor Decomposition | 0.877 ± 0.073 | 0.938 ± 0.053 | 0.888 ± 0.067 |
| | Mean | Concatenation | 0.832 ± 0.038 | 0.921 ± 0.018 | 0.858 ± 0.026 |
| | | Expression Only | 0.863 ± 0.016 | **0.94 ± 0.027** | 0.888 ± 0.014 |
| | | Tensor Decomposition | 0.803 ± 0.065 | 0.903 ± 0.053 | 0.809 ± 0.079 |

**Supplementary Table 3. SNOWFLAKE pipeline architecture search for graph layer, graph pooling layer, and morphology fusion type.**

**Reference**

1. S. Black, D. Phillips, J. W. Hickey, J. Kennedy-Darling, V. G. Venkataraaman, N. Samusik, Y. Goltsev, C. M. Schürch, G. P. Nolan, CODEX multiplexed tissue imaging with DNA-conjugated antibodies. *Nat. Protoc.* **16**, 3802–3835 (2021).

2. R. Rashid, G. Gaglia, Y.-A. Chen, J.-R. Lin, Z. Du, Z. Maliga, D. Schapiro, C. Yapp, J. Muhlich, A. Sokolov, P. Sorger, S. Santagata, Highly multiplexed immunofluorescence images and single-cell data of immune markers in tonsil and lung cancer. *Sci. Data*. **6**, 323 (2019).

3. D. Schapiro, A. Sokolov, C. Yapp, Y.-A. Chen, J. L. Muhlich, J. Hess, A. L. Creason, A. J. Nirmal, G. J. Baker, M. K. Nariya, J.-R. Lin, Z. Maliga, C. A. Jacobson, M. W. Hodgman, J. Ruokonen, S. L. Farhi, D. Abbondanza, E. T. McKinley, D. Persson, C. Betts, S. Sivagnanam, A. Regev, J. Goecks, R. J. Coffey, L. M. Coussens, S. Santagata, P. K. Sorger, MCMICRO: a scalable, modular image-processing pipeline for multiplexed tissue imaging. *Nat. Methods*. **19**, 311–315 (2022).

4. Q. Xu, P. Milanez-Almeida, A. J. Martins, A. J. Radtke, K. B. Hoehn, C. Oguz, J. Chen, C. Liu, J. Tang, G. Grubbs, S. Stein, S. Ramelli, J. Kabat, H. Behzadpour, M. Karkanitsa, J. Spathies, H. Kalish, L. Kardava, M. Kirby, F. Cheung, S. Preite, P. C. Duncker, M. M. Kitakule, N. Romero, D. Preciado, L. Gitman, G. Koroleva, G. Smith, A. Shaffer, I. T. McBain, P. J. McGuire, S. Pittaluga, R. N. Germain, R. Apps, D. M. Schwartz, K. Sadtler, S. Moir, D. S. Chertow, S. H. Kleinstein, S. Khurana, J. S. Tsang, P. Mudd, P. L. Schwartzberg, K. Manthiram, Adaptive immune responses to SARS-CoV-2 persist in the pharyngeal lymphoid tissue of children. *Nat. Immunol.* **24**, 186–199 (2023).

5. J.-R. Lin, B. Izar, S. Wang, C. Yapp, S. Mei, P. M. Shah, S. Santagata, P. K. Sorger, Highly multiplexed immunofluorescence imaging of human tissues and tumors using t-CyCIF and conventional optical microscopes. *eLife*. **7**, e31657 (2018).
